# Dlk1-Dio3 cluster miRNAs regulate mitochondrial functions in Duchenne muscular dystrophy

**DOI:** 10.1101/2021.10.20.464950

**Authors:** Ai Vu Hong, Nathalie Bourg, Peggy Sanatine, Jerome Poupiot, Karine Charton, Evelyne Gicquel, Emmanuelle Massourides, Marco Spinazzi, Isabelle Richard, David Israeli

## Abstract

**Background:** Duchenne Muscular Dystrophy (DMD) is a severe muscle disease caused by impaired expression of dystrophin. While mitochondrial dysfunction is thought to play an important role in DMD, the mechanism of this dysfunction remains to be clarified. We recently identified in DMD and in other muscular dystrophies the upregulation of a large number of the Dlk1-Dio3 clustered miRNAs (DD-miRNAs), in both the muscle and the serum. The objective of the present study was to define the biological functions of DD-miRNAs in skeletal muscle, particularly in the context of muscular dystrophy.

**Methods:** DD-miRNAs expression pattern was characterized *in vitro* and *in vivo*, in normal and dystrophic situations. Epigenomic characterization was performed, to elucidate the molecular control of DD-miRNAs dysregulation. The biological effect of muscle DD-miRNAs dysregulation was investigated by an *in vivo* simultaneous **overexpression** of 14 DD-miRNAs in the wild-type muscle, together with CRISPR-Cas9-based **knockdown** of the entire DD-miRNA cluster in an iPS-derived myotubes. Omics data and bioinformatics tools were used for the prediction of DD-miRNAs biological functions, and functional characterization of mitochondrial pathways was performed.

**Results:** We found that DD-miRNAs dysregulation is not specific to DMD since observed in mouse models for other muscular dystrophies. We showed that DD-miRNAs expression in mdx, is reduced in satellite cells, but highly upregulated in regenerating myofibers, suggesting a myofibers origin of DD6miRNA upregulation in muscular dystrophy in both muscles and serum. We demonstrated that upregulation of DD-miRNAs in the dystrophic muscle is controlled epigenetically by DNA and histone methylation (p<0.0001 and p=0.001, respectively) at the Intergenic Differentially Methylated Region (IG-DMR) of Dlk1-Dio3 locus. Transcriptomic analysis revealed a substantial overlap between the dystrophic muscle of the mdx mouse and the normal muscle that overexpressed 14 DD-miRNAs. Bioinformatics analysis predicted that DD-miRNAs could regulate mitochondrial functions. The ectopic overexpression of 14 DD-miRNAs, in the healthy muscle, resulted in a drastic downregulation of mitochondrial oxidative phosphorylation (OxPhos) (NES=-2.8, p=8.7E-17), similarly to the level in dystrophic muscles of mdx mice and DMD patients (NES=-2.88, p=7.7E-28). Knocking down the entire DD-miRNA cluster in iPS-derived myotubes resulted in increased mitochondrial OxPhos expression and activities.

**Conclusions:** The present study provides evidence for the modulation of mitochondrial activity in the dystrophic muscle by the upregulated DD-miRNAs and supports an updated model for mitochondrial dysfunction in DMD. The regulation of mitochondrial OxPhos by DD-miRNAs may have a broader impact beyond DMD in physiological and pathological situations of muscle adaptation and regeneration.

## INTRODUCTION

Duchenne Muscular Dystrophy (DMD) is an X-linked severe progressive muscle disease caused by mutations in the *Dmd* gene that codes for the dystrophin protein. The disease severely affects the motor function and leads to the premature death of the patient, primarily due to respiratory and cardiac failures ^1^. In striated muscle, dystrophin and its associated proteins link the intracellular cytoskeletal network and the extracellular matrix to stabilize myofibers. The lack of dystrophin protein results in a structural defect in the sarcolemma and impairs contractile activity. The lack of dystrophin promotes a pathological cascade, that includes calcium overload, hyperactive proteases, oxidative stress, mitochondrial dysfunction and damages, chronic inflammation, muscle degeneration and impaired regenerative capacity, and in advanced stage the replacement of the contractile tissue by fibrotic and fat tissues ^1,2^.

We recently identified a coordinated dysregulation in DMD of many miRNAs originated from Dlk1-Dio3 locus ^3-5^. The imprinted Dlk1-Dio3 locus hosts the largest miRNA mega-cluster in the human genome, in addition to other non-coding RNAs and three protein-coding genes (*Dlk1*, *Rtl1*, and *Dio3*) and is highly conserved in the mammalian genomes ^6^. Dlk1-Dio3 miRNAs (DD-miRNAs) were shown to play a critical role in fetal development and postnatal growth ^6-8^. Initial indications for DLK1-DIO3 locus involvement in the muscular system came from the identification of the muscular hypertrophy Callipyge phenotype in the sheep^9^. Consistently, patients carrying a genetic defect in Dlk1-Dio3 locus present hypotonia and muscle metabolic deficiencies. However, the biological functions of these miRNAs in the context of muscular dystrophy remains relatively unexplored.

In the present study, we showed that the DD-miRNA dysregulation is associated to muscle regeneration since, in addition to DMD, it was found also in several mouse models for other muscular dystrophies, as well as in regenerating normal muscle. We used the mdx mouse model for the investigation of DD-miRNAs expression control and biological functions in DMD. We found that in the dystrophic muscle the upregulation of DD-miRNAs correlates with epigenetic changes at two regulatory regions of the Dlk1-Dio3 locus. Combined analysis of DD-miRNAs target prediction, and of the transcriptome of dystrophic muscles, suggested that DD-miRNAs may target mitochondrial metabolism, and particularly the oxidative phosphorylation (OxPhos) system. Indeed, *in vivo* overexpression of 14 selected DD-miRNAs, in healthy muscles, drastically reduced mitochondrial OxPhos, and importantly, partly resembled the transcriptome of the dystrophic muscle. Furthermore, knocking down the entire DD-miRNAs cluster in hiPSC-derived myotubes resulted in increased RNA and proteins expression of OxPhos components, and the increased activity of the mitochondrial 1-5 complexes. Our data suggesting a cooperative regulation by DD-miRNAs of mitochondrial functions in DMD and possibly in other situations of muscle regeneration.

## MATERIALS AND METHODS

### Animal care and use

All animals were handled according to French and European guidelines for human care and the use of experimental animals. All procedures on animals were approved by Genethon’s ethics committee under the numbers CE10-122, CE10-123, CE10-124, CE10-127, and CE12-039. C57Bl10, C57Bl6, and C57BL/10ScSn-Dmd^mdx^/J mice were obtained from Charles River laboratories. Mice were housed in a SPF barrier facility with 12-h light, 12-h dark cycles, and were provided with food and water ad libitum. Only male mice were used in the present study. The animals were anesthetized with a mix of ketamine (100 mg/kg) and xylazine (10 mg/kg), or with isoflurane, for blood samples. For intramuscular injections, a volume of 25 µl containing 1.0E10vg AAV vectors was injected into each TA muscle. For the duration of the study, all animals were observed at least once a day. All animals were weighed on the day of treatment as well as on the day of the necropsy.

### AAV vectors expressing multiple miRNAs

The 14 pre-miRNA sequences were obtained from UCSC, spanning 100 nucleotides before and after the mature miRNA sequences. The pre-miRNA sequences were then arranged consecutively which respect the genomic sequences of the miRNAs. Two or three pre-miRNAs were used per AAV construct.

### Transcriptomic analysis

Differentially expressed genes were identified by DESeq2 R package. Pathway analysis was performed in R-Studio, either by over-representation methods using Gene Ontology and ReactomePA or functional class scoring using Geneset Enrichment Analysis (GSEA).

### Computational prediction of DD-miRNA targets

Targets of Dlk1-Dio3 miRNAs were predicted by using R package miRNAtap ^10^, compiling data from five public databases: DIANA, Targetscan, PicTar, Miranda, and miRDB. The targets used in subsequent analysis are genes predicted by at least two databases.

### Genomic deletion using the CRISPR/Cas9 and screening for bi-allelic deletion clones

Different combinations of single-guide RNAs (sgRNAs) targeting two ends of IG-DMR region were tested in 911 human cells for cutting efficacy. Selected sgRNAs were then used to delete IG-DMR region in hiPS cells, followed by clonal selection and myogenic differentiation.

### Data presentation and statistical analysis

The results were presented as mean ± standard error of the mean (SEM) of at least three replicates. PRISM 7.01 program (GraphPad) and R-Studio (version 4.0.3) were used for statistics. Comparisons between two groups were done using student t-test. Comparisons between two groups according to the levels of two categorical variables were done using two-way ANOVA. Significance was defined as *p<0.05, **p<0.01, ***p<0.001.

Other material and methods are provided in the **extended materials and methods** in the supplemental material.

## RESULTS

### Upregulation of DD-miRNA in regenerating myofibers and in the serum is associated to muscle regeneration

In previous studies, we identified a dysregulation of a large number of miRNAs of the Dlk1-Dio3 cluster (DD-miRNAs) in the serum of the GRMD dog, a model for DMD ^4^, and in the plasma of DMD patients ^3^. In the GRMD model, we found that DD-miRNAs are among the most highly upregulated miRNA, not only in the circulation, but also in the muscle ^5^. In the present study, we confirmed this observation across species. In particular, we quantified DD-miRNAs in muscle biopsies of DMD patients (**Figure 1A**, n=4), and mdx mouse (**Figure 1B**, 5-week-old mice, diaphragm muscle, n=6) and confirmed their upregulation in the dystrophic muscle. Thus, DD-miRNAs are upregulated in DMD in the muscles and in the serum.

**Figure 1:**
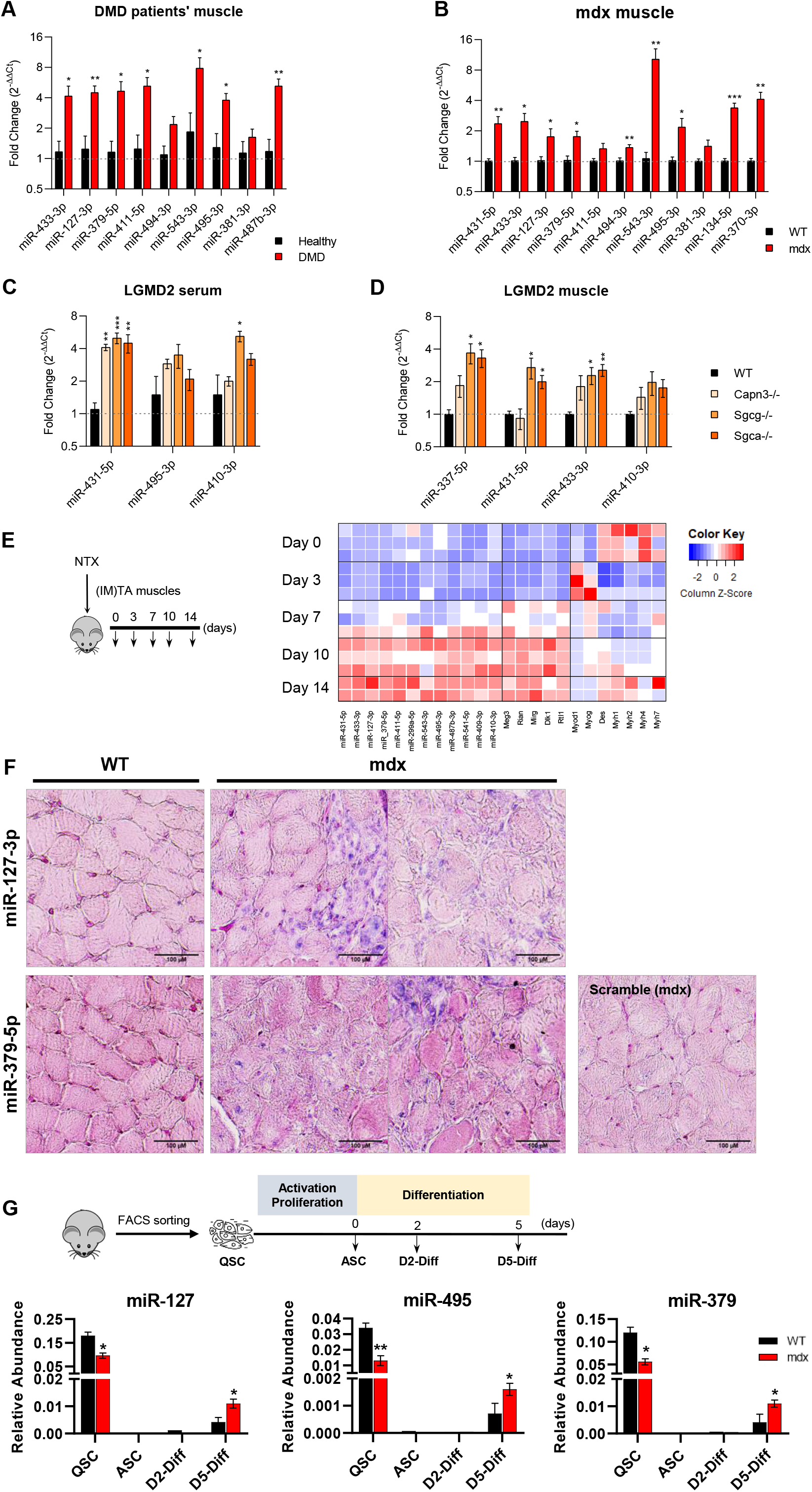
Characterization of DD-miRNAs dysregulation in the regenerating muscle. **A**. Relative expression levels of DD-miRNAs in muscle biopsies of DMD patients compared to healthy controls (n=4). **B**. Relative expression levels of DD-miRNAs in diaphragm muscle of mdx mouse model compared to C57Bl10 control (n=6). **C**-**D**. Relative expression levels of DD-miRNAs in the serum (C) and muscle (D) of 3 LGMD mouse models compared to C57Bl6 control (n=4). **E**. Heatmap presenting expression level of DD-miRNAs, other transcripts of the Dlk1-Dio3 locus, and myogenic transcription factors during *in vivo* acute muscle regeneration (n=2-3). The relative expression level is illustrated by column Z-scores, colored from blue to red, indicating from lowest to highest expression. **F.** ISH of miR-127-3p (upper panels) and miR-379-5p (lower panels) in diaphragm of mdx and controls. Nucleus was colored in pink while miRNAs of interest was colored in dark blue. Scramble probes served as negative control. Scale bar: 100 μm. **G.** Relative expression of three DD-miRNAs through *in vitro* differentiation of satellite cells from muscles of 5-week-old mdx and C57Bl10 control mice (n=3). QSC: quiescent satellite cells; ASC: activated satellite cells; D2/D5-Diff: Day 2/5 differentiated myotubes. IM: intramuscular injection. Data in all graphical presentations are presented as mean±SEM. Statistics were performed with Student t-test. *P < 0.05; **P < 0.01; ***P < 0.001.

To define whether these dysregulations are specific to DMD, we quantified DD-miRNAs in serum and muscle biopsies of a collection of mouse models for limb-girdle muscular dystrophies, including LGMDR1 (calpainopathy), LGMDR5 (gamma sarcoglycanopathy) and LGMDR3 (alpha sarcoglycanopathy). Upregulation of DD-miRNAs was identified in both serum and muscles of all these disease models (n=4, **Figure 1C-D**) and, interestingly, in a severity dependent manner. DD-miRNAs are therefore expressed in the healthy muscle and upregulated further in distinct muscular dystrophies. Next, we analyzed the expression of DD-miRNAs during muscle regeneration in the model of myotoxin-induced injury. The levels of a collection of DD-miRNAs, along with other maternal and paternal transcripts from the same locus (*Meg3*, *Rian*, *Mirg*, *Dlk1*, and *Rtl1*), increased gradually from day 7 to 14 post-injury, confirming upregulation in the regenerating muscle independently of a genetic defect **(Figure 1E)**.

In addition to the muscle, the postnatal expression of DD-miRNAs was shown also in the brain ^11,12^. In agreement, we found the highest expression DD-miRNAs in the brain, which was followed by the skeletal muscle, and a much lower levels in all other tissues and organs. However, the upregulation of DD-miRNAs in muscular dystrophy was found in the mdx mouse only in the skeletal muscle **(Supplemental Figure 1A).** Thus, serum upregulation of DD-miRNAs in this mouse model is likely originating from the muscle. Since the skeletal muscle is a complex tissue composed of several cell types, we attempted to clarify which are the specific sites and cells that are expressing the DD-miRNAs within the muscle. We performed *in situ* hybridization (ISH) of two relatively highly expressed DD-miRNAs, miR-127-3p and miR-379-5p, in the diaphragm of 5-week-old mdx mice and their respective controls. Both miRNAs presented similar pattern of high expression in muscle areas of intensive muscle regeneration, composed of small-diameter centranucleated regenerating myotubes and of muscle mono-nucleated cells (MMNC) (**Figure 1F**), thus both MMNC and regenerating myotubes may contribute to the upregulation of DD-miRNAs in the serum.

To distinguish between these two possibilities (for the origin of upregulation in the dystrophic muscle), we profiled DD-miRNAs in FACS-sorted MMNC subpopulations, and *in vitro* in differentiated myotubes. DD-miRNAs expression was not upregulated in any of the mdx-derived MMNC fractions. On the contrary, in agreement with ^13^, DD-expression was repressed in the freshly-sorted satellite cells (QSC) **(Supplemental figure 1B)**. To test DD-miRNAs expressions in myotubes, satellite cells from hind-limbs muscles of mdx and control mice were FACS-sorted, grown, and differentiated *in vitro* for 5 days. Profiling of three representative DD-miRNAs confirmed their reduced level in mdx QSC. DD-miRNAs expression then dropped down sharply at the beginning of the differentiation stage. Expression however raised up in day-5 differentiated myotubes, to a level significantly higher in mdx than in controls (**Figure 1G**, n=3). Thus, muscle upregulation of DD-miRNAs is seemingly coming from regenerating myofibers and not from the MMNC. In summary, the postnatal expression of DD-miRNAs in the mouse is the highest in brain, followed by skeletal muscle. In various muscular dystrophies, DD-miRNAs expression is upregulated both in muscle and serum. In the muscle, the upregulation is occuring in the regenerating fibers. The data could also suggest that serum upregulation of DD-miRNAs is linked to their increased expression in regenerating myotubes, likely through an active mechanism rather than passive leakage through damaged sarcolemma, since these regenerated fibers are not in the process of degeneration.

### Epigenetic modifications control DD-miRNAs upregulation in mdx muscle

We then investigated the upstream regulation of the DD locus. Two central regulatory regions are known to control the expression of maternal transcripts of the Dlk1-Dio3 imprinted locus: the Intergenic Differentially Methylated Region (IG-DMR) and the Meg3 Differentially Methylated Region (Meg3-DMR), which are known to be subjected to histone modifications and DNA methylation ^6,14^. First, we used the C2C12 myoblast cell line in order to test if the Dlk1-Dio3 locus is epigenetically controlled in the myogenic linage. C2C12 cells were incubated with 5-Azacytidine, a DNA methylation inhibitor, and with Givinostat and TSA, both being histone deacetylation (HDAC) inhibitors. The inhibition of DNA methylation and of HDAC activity resulted in significant increased expression of genes of the Dlk1-Dio3 locus (**Supplemental Figure 2A-C**), indicating that both types of epigenetic modifications participate in the control of gene expression in the Dlk1-Dio3 locus in the myogenic lineage.

We then asked if epigenetic changes of the mdx diaphragm are occurring specifically at the Dlk1-Dio3 locus. Bisulfite sequencing demonstrated a reduction in DNA methylation level at IG-DMR (**Figure 2A**, p=1.06E-05, two-way ANOVA test), but not Meg3-DMR. Chromatin immuno-precipitation (ChIP) followed by qPCR was used for the detection of histone modifications of IG-DMR and Meg3-DMR regulatory regions. We found a significantly reduced percentage of H3K27me3 histone repressive marker in both IG-DMR (p=0.001, two-way ANOVA test) and Meg3-DMR (p=0.009, two-way ANOVA test) in the mdx diaphragm compared to control **(Figure 2B**).

**Figure 2:**
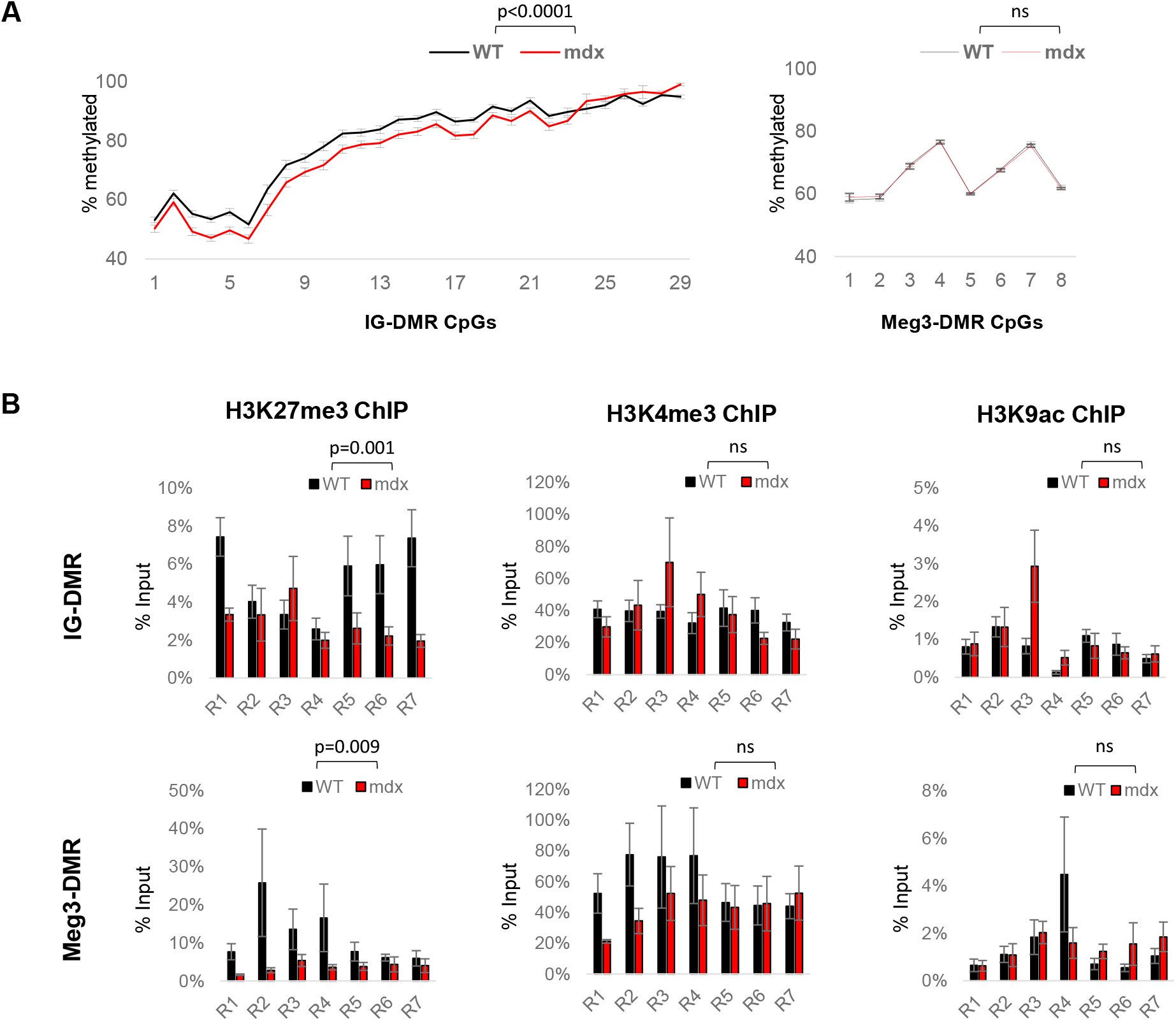
DD-miRNAs expression is epigenetically controlled in DMD. **A**. DNA methylation levels of 29 CpGs in IG-DMR and 8 CpGs in Meg3-DMR. **B**. ChIP-qPCR of H3K27me3, H3K4me3, and H3K9ac in IG-DMR and Meg3-DMR. The PCR amplicons are shown schematically in **Supplemental Figure 2E**. Statistics were performed by 2-way ANOVA test. Data are presented as mean±SEM. All experiments were performed in the diaphragms of 5-week-old mdx and C57Bl10 controls.*P < 0.05; **P < 0.01; ***P < 0.001.

### *In vivo* DD-miRNAs overexpression reduces muscle mass

To investigate the functions of DD-miRNAs *in vivo* in skeletal muscle and since miRNAs often act synergistically, we therefore overexpressed simultaneously a selection of miRNAs in the wild-type muscle using recombinant Adeno-Associated Virus (AAV-derived vectors). We selected 14 miRNAs based on their expression and dysregulation levels in dystrophic serum, skeletal muscle and quiescent satellite cells **(Supplemental Figure 3A, Supplemental Table 4, Figure 3A)**.

**Figure 3:**
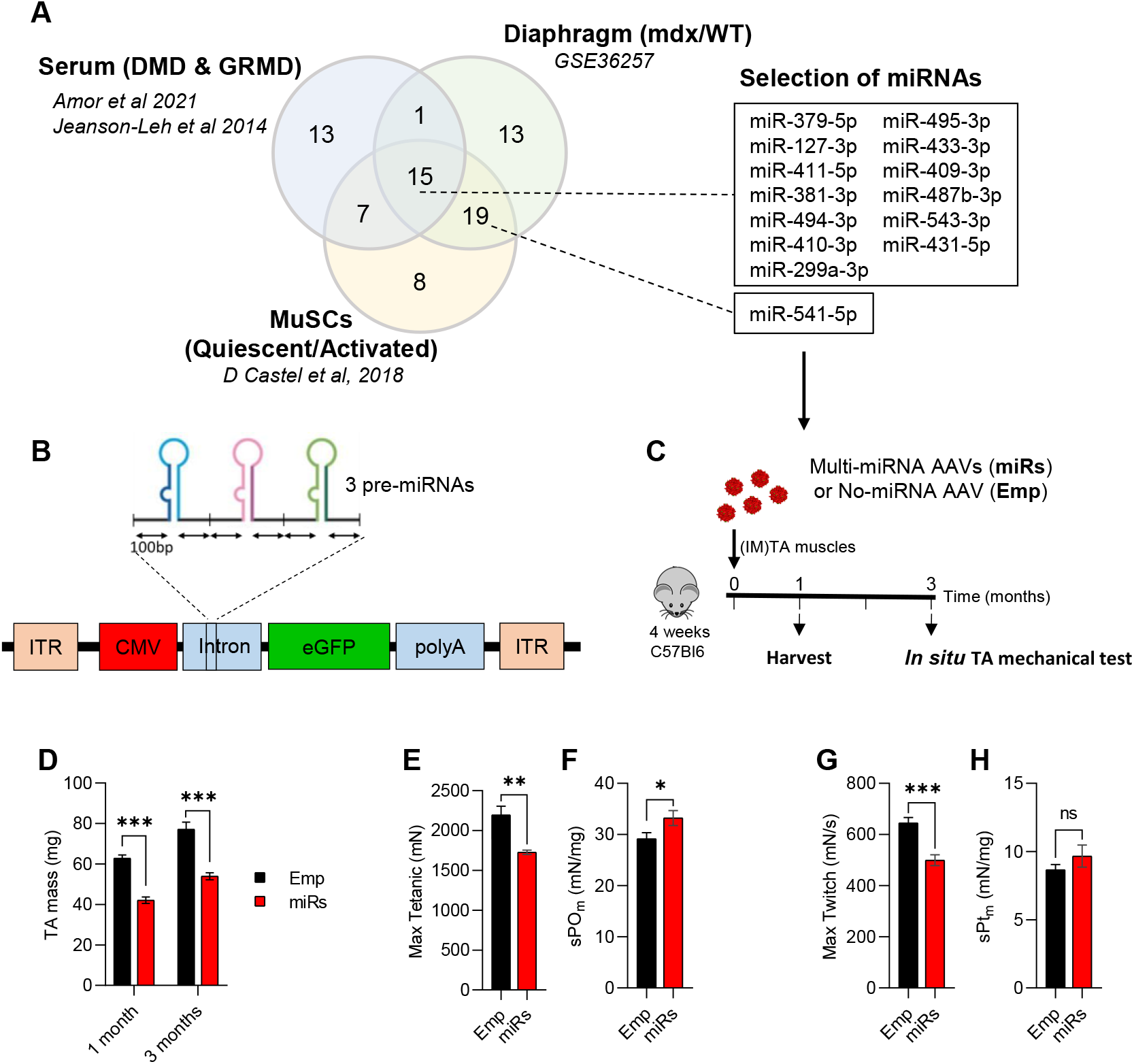
Selection of DD-miRNAs and their *in vivo* overexpression by AAV vectors. **A.** Of the selected 14 DD-miRNAs, 13 are commonly dysregulated in the serum of DMD patients and GRMD dog models ^3,4^, in the diaphragm muscle of the mdx mouse (GEO accession: GSE3625 **B**. AAV construction strategy for the overexpression of 14 DD-miRNAs. **C**. Experimental design of the *in vivo* overexpression of DD-miRNAs by the intramuscular co-injection of 5 AAV vectors to the TA muscle of a wild-type 4-week-old mouse (miRs). Injection of AAV vector with same expression cassette but pre-miRNAs sequences at the same titer (Emp) served as control in the experiment. **D**. Masses of TA muscles 1-month and 3-month post-injection (n=10). **E-H.** I*n situ* max tetanic (**E**) and max twitch (**G**) forces of treated muscles 3-month post-injection (n=5), and its normalization to muscle masses (**F**, **H**). Data are presented as mean±SEM. Statistics were performed with Student t-test. *P < 0.05; **P < 0.01; ***P < 0.001.

The mouse pre-miRNA sequences of these 14 miRNAs were sub-cloned into five AAV expression vectors (four that contained each a tandem of three DD-miRNAs and one with two DD-miRNAs) that co-express the GFP protein, as schematically described in (**Figure 3B**). These five AAV vectors were co-administrated by an intramuscular (IM) injection into the tibialis anterior (TA) muscle of 4-week-old C57Bl6 control mice. An AAV vector with the same expression cassette but devoid of pre-miRNA sequences served as negative control. The protocol included two time points (1 and 3 months post-injection) with a muscle function tests at the 3-month group **(Figure 3C)**. Analysis of the injected muscles confirmed the overexpression of all injected DD-miRNAs (**Supplemental Figure 3B**) to a comparable level to endogenous DD-miRNAs in the diaphragms of the mdx mice and of DMD muscle biopsies (**Supplemental Figure 3C**). DD-miRNAs overexpressing TA muscles presented a significantly reduced muscle mass at both 1 and 3 months after injection (49.13% and 43.15%, respectively; **Figure 3D)**. At 3 months post injection, *in situ* mechanical force measurement showed that both absolute value of maximum tetanic force and maximum twitch force of DD-miRNA overexpressed muscles significantly dropped at 27.2% and 29.1%, respectively (**Figure 3E, G**, n=5). When reported to the muscle mass, the normalized values of tetanic force and twitch force were slightly higher in the presence of DD-miRNAs (**Figure 3F, H**, n=5), indicating that the reduction in mechanical force was due to the loss of muscle mass.

### The transcriptomic changes following ectopic overexpression of 14 DD-miRNAs overlap with the dysregulation in mdx dystrophic muscle

To understand to which extend the dysregulation of DD-miRNAs contributes to the transcriptomic changes of the dystrophic muscle, we compared the transcriptomic profile of the dystrophic muscles to that of the healthy muscles overexpressing DD-miRNAs. In the dystrophic diaphragm, 8309 differentially expressed genes (DEGs) were identified (n=4, BH-adjusted p-value < 0.05) (**Figure 4A, C**). In one-month treated muscles injected with AAV-DD-miRNAs, DD-miRNAs overexpression resulted in 2048 DEGs (n=3, BH-adjusted p-value < 0.05) (**Figure 4A, D**), of which 74.2% (1520 out of 2048) were also dysregulated in the dystrophic muscle (**Figure 4A**). Furthermore, a highly significant correlation was observed between the level of dysregulation of these 1520 common DEGs within the two datasets (Pearson correlation test, R=0.73, p-value < 2.2E-16) (**Figure 4B**). These data indicate that the majority of changes driven by DD-miRNAs overexpression in the normal muscle are included in the transcriptomic dysregulation seen in the mdx muscle. Of note, we found that many of the highly correlating repressed transcripts are of mitochondrial genes **(Cyan dots in Figure 4B)**.

**Figure 4:**
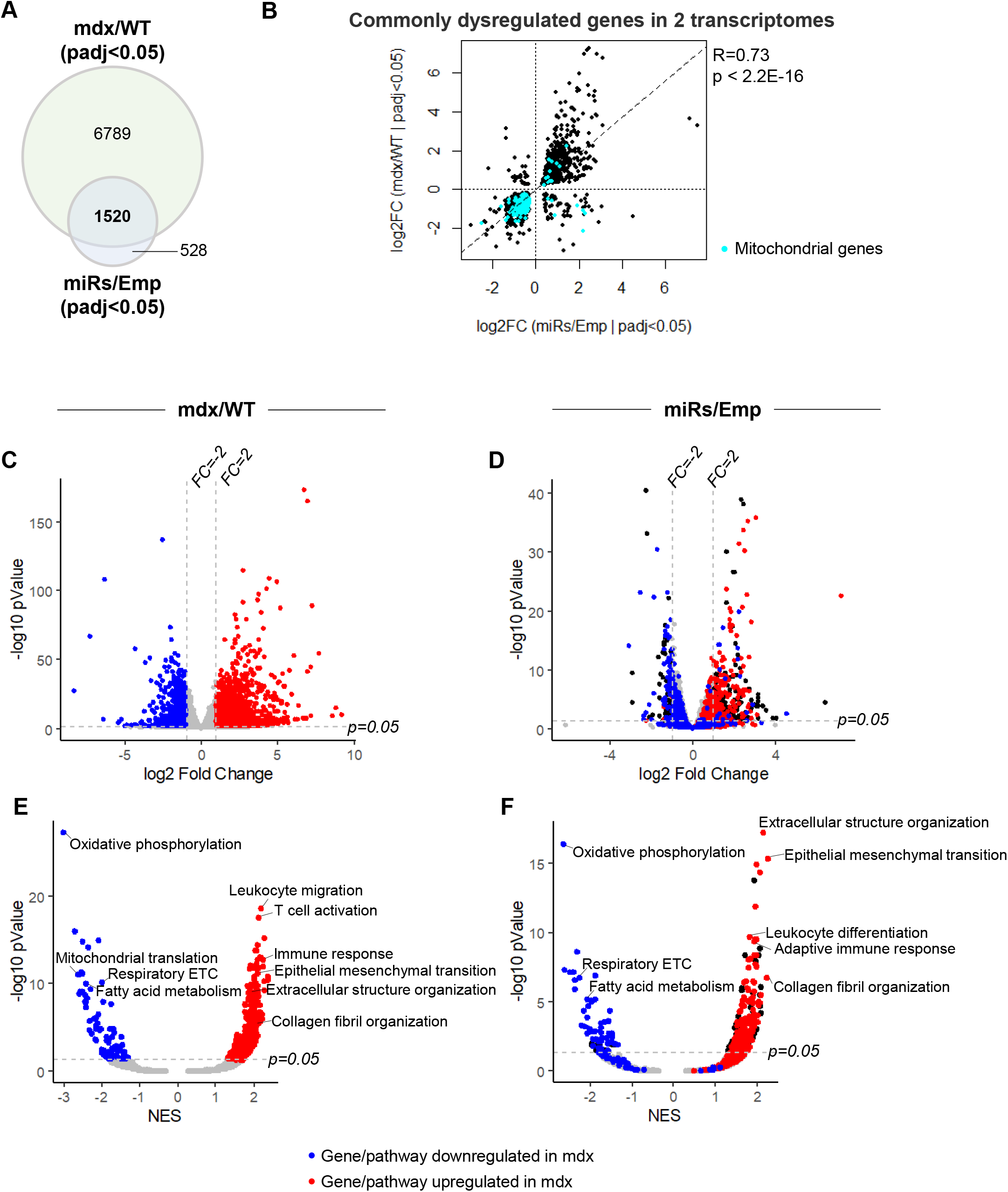
Comparison of the transcriptomic profiles between 14 DD-miRNAs ectopic expression and the mdx dystrophic muscle. **A.** Venn diagram of dysregulated genes in mdx diaphragm compared to wild-type control and in DD-miRNAs-overexpressed TA muscle compared to AAV control. **B**. Dot plot of log2 fold change values of 1520 commonly dysregulated genes in the 2 transcriptomes and its Pearson correlation test. Cyan dots represent mitochondrial-related genes (Mitocarta 2.0). **C, D**. Volcano plots of mdx/WT transcriptome (**C**) and miRs/Emp transcriptome (**D**). Blue dots represent significantly down-regulated genes with fold change of mdx versus WT less than −2 while red dots represent significantly up-regulated genes with fold change of mdx versus WT greater than 2 (BH-adjusted p-value < 0.05). **E, F**. Volcano plots of all Hallmark and Gene Ontology (Biological Process) gene sets from GSEA analysis, comparison of mdx/WT transcriptome (**E**) and miRs/Emp transcriptome (**F**). Blue dots represent significantly down-regulated pathways in mdx compared to WT while red dots represent significantly up-regulated pathways in mdx compared to WT (BH-adjusted p-value < 0.05).

Next, gene set enrichment analysis (GSEA) was utilized to interpret the transcriptomic changes at the pathway level ^15^. As expected, pathways related to immune response and fibrosis progression were found highly upregulated in the mdx muscle, and many metabolic pathways were significantly downregulated (**Figure 4E**). Strikingly, high-level overlap of pathway dysregulation was identified in the DD-miRNAs overexpressed muscle, compared to negative controls (**Figure 4F; Supplemental Table 5**). Consistent with the down-regulation of mitochondrial transcripts, the pathway analysis revealed downregulation of mitochondrial pathways and particularly of oxidative phosphorylation, which was by far the most downregulated pathway in both systems (the dystrophic muscle and the healthy muscles overexpressing DD-miRNAs).

Taken together, the data supported that about 75% of the transcripts which are potentially repressed by DD-miRNAs, may participate in significant gene regulation in the dystrophic diaphragm of the mdx mouse, which potentially account to up to approximately 20% of the diaphragm transcriptome dysregulation.

### Dlk1-Dio3 miRNAs affect metabolic pathways in the skeletal muscle

Next, we employed a bioinformatics approach to identify potential 14 selected DD-miRNAs’ target genes. First, five prediction tools (DIANA, Targetscan, PicTar, Miranda, and miRDB) were used, generating 5295 candidate targets predicted by at least two different tools **(Figure 5A)**. The list of these predicted targets was subsequently crossed with the lists of genes downregulated in DD-miRNAs injected muscles and in the mdx diaphragm (that naturally overexpresses the DD-miRNAs). The analysis resulted in 269 DD-miRNAs predicted targets **(Supplemental Table 6, Figure 5A)**. These 269 genes were subjected to gene ontology (GO) classification, which identified metabolism-related terms at the highest p-values, including mTOR signaling, pyruvate metabolism, TCA cycle and mitochondrial electron transport chain **(Figure 5B)**. Downregulation of a selection of these genes was then experimentally validated in the dystrophic muscle (5-week-old mice, diaphragm muscle, n=4) and AAV-DD-miRNAs-injected TA muscles (n=4) at the levels of mRNA (**Figure 5C**) and protein (**Figure 5D**). These data strongly support that DD-miRNAs are involved in the metabolic adaptive response of the dystrophic muscle.

**Figure 5:**
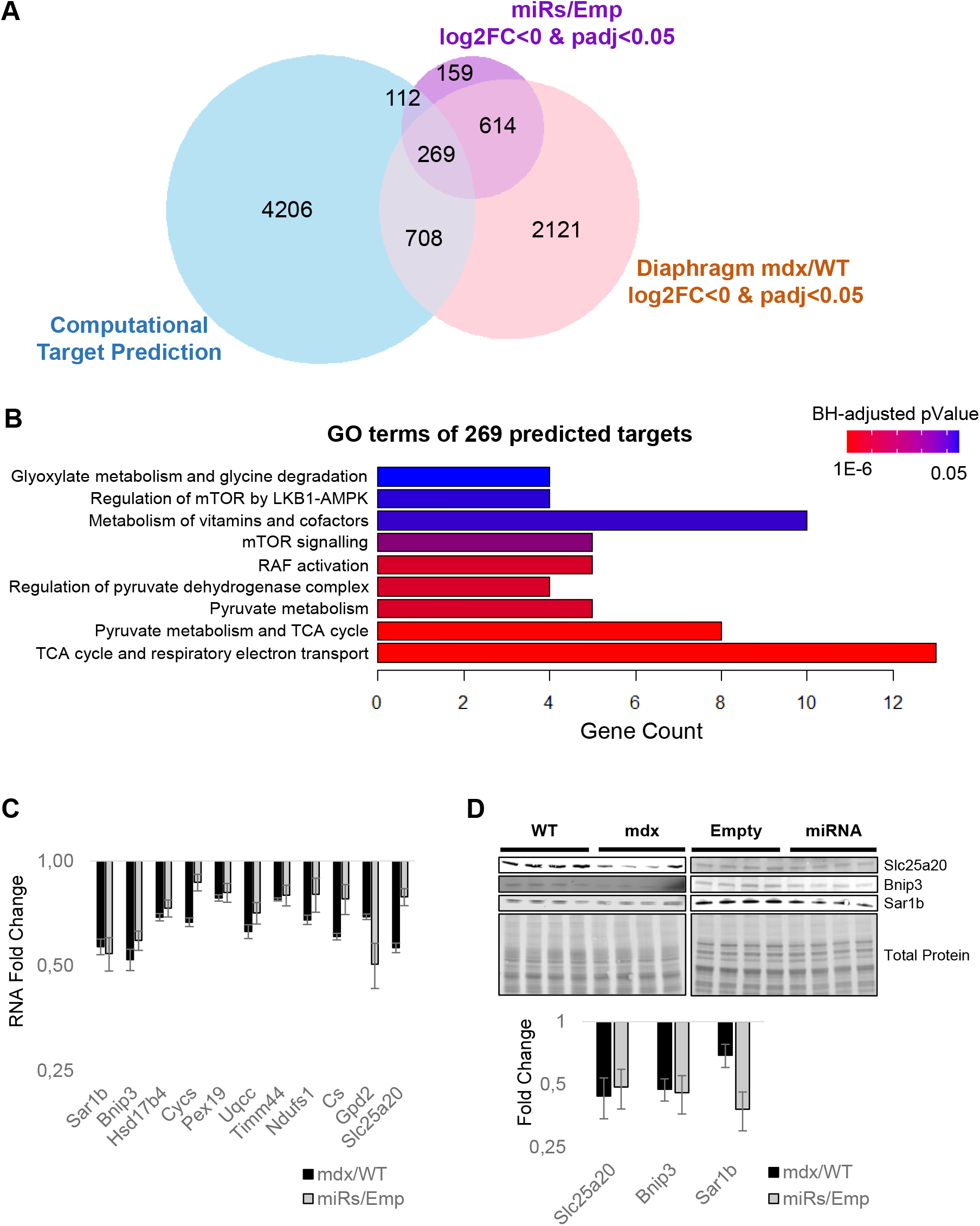
DD- miRNAs are predicted to target metabolic pathways in the skeletal muscle. **A.** Venn diagram of significantly down-regulated genes in mdx/WT, of miRs/Emp transcriptome and of DD-miRNAs’s predicted target genes. **B**. Pathway analysis of the 269 predicted targets for DD-miRNAs identified in **A**. **C-D**. Validation at RNA (**C**) and protein (**D**) levels of selected predicted targets (n=4). Data in **C** and **D** are presented as mean±SEM. Statistics were performed with Student t-test. *P < 0.05; **P < 0.01; ***P < 0.001.

### DD-miRNAs affect mitochondrial functions in the dystrophic muscle

Clustered miRNAs are thought to coordinately regulate genes that are functionally related ^16^. In the dystrophic muscle, we have recently shown that one specific DD-miRNA, namely miR-379-5p, is acting on the mitochondrial oxidative phosphorylation pathway ^5^. Taken together, we hypothesized that, in the dystrophic muscle, DD-miRNAs might coordinately regulate mitochondrial functions. In order to estimate the global level of mitochondrial adaptation that occurs in the dystrophic muscle, we used our transcriptomic data of the mdx diaphragm and of the DD-miRNAs overexpressed muscles, combined with the Mitocarta database of mitochondrial genes ^17^. Of the 7406 dysregulated (adjusted p-value < 0.05) transcripts in the mdx diaphragm, 788 (10.64%) were found to be mitochondrial. In order to estimate the relative role DD-miRNAs in this mitochondrial adaptive response, we found that of the 2048 dysregulated transcripts in the TA-DD-miRNAs, 265 (12.94%) were found to be mitochondrial **(Supplemental Figure 4A)**, of which 250 (out of 265) were commonly dysregulated in the mdx diaphragm and therefore potentially regulated by DD-miRNAs in the mdx diaphragm. Thus, of the 788 mitochondrial transcripts dysregulated in the mdx diaphragm, about one-third could be impacted by DD-miRNAs. This is however an underestimation, because only 14 of all dysregulated DD-miRNAs ^3,4^, were ectopically overexpressed in the TA muscle. We then attempted to characterize in details the mitochondrial effect of the DD-miRNAs overexpressed muscle as compared to the mdx diaphragm. Mitochondrial DNA copy number **(Figure 6A, D)**, citrate synthase levels (**Figure 6B, E**), and resting ATP concentration level (**Figure 6C, F**) were both significantly reduced in mdx compared to WT and in TA-DD-miRNAs compared to TA-Empty control. Immunohistochemistry of cytochrome oxidase (COX) and succinic dehydrogenase (SDH) in mdx diaphragm and TA-DD-miRNAs muscle confirmed a reduction of mitochondrial activities in those two muscles (**Figure 6G-H**). Taken together, the data supports that DD-miRNAs are acting on mitochondrial metabolism in dystrophic muscle. The gene expression and pathway analyses described above indicated that mitochondrial oxidative phosphorylation might be one of the most affected mitochondrial functions downstream to the dysregulation of DD-miRNAs (**Figure 4E-F, Supplemental Figure 4B-C**). We therefore attempted to characterize it further. Similar reduction of OxPhos RNA level was observed in muscle biopsies from two independent large cohorts of young DMD patients compared to its age-matched healthy controls (**Supplemental Figure 4E, G**) (data was taken from GEO accession: GSE6011 ^18^ and GEO accession: GSE38417). As expected, expression level of Meg3 transcripts are highly significantly upregulated within the DMD groups’ muscles (GSE6011: n=13/23, p-value=0.0009; GSE38417: n=6/16, p-value<0.0001) (**Supplemental Figure 4D, F**). This data demonstrates a reduced OxPhos and its negative correlation with Dlk1-Dio3 maternal transcripts in muscle biopsies of human DMD patients too.

**Figure 6:**
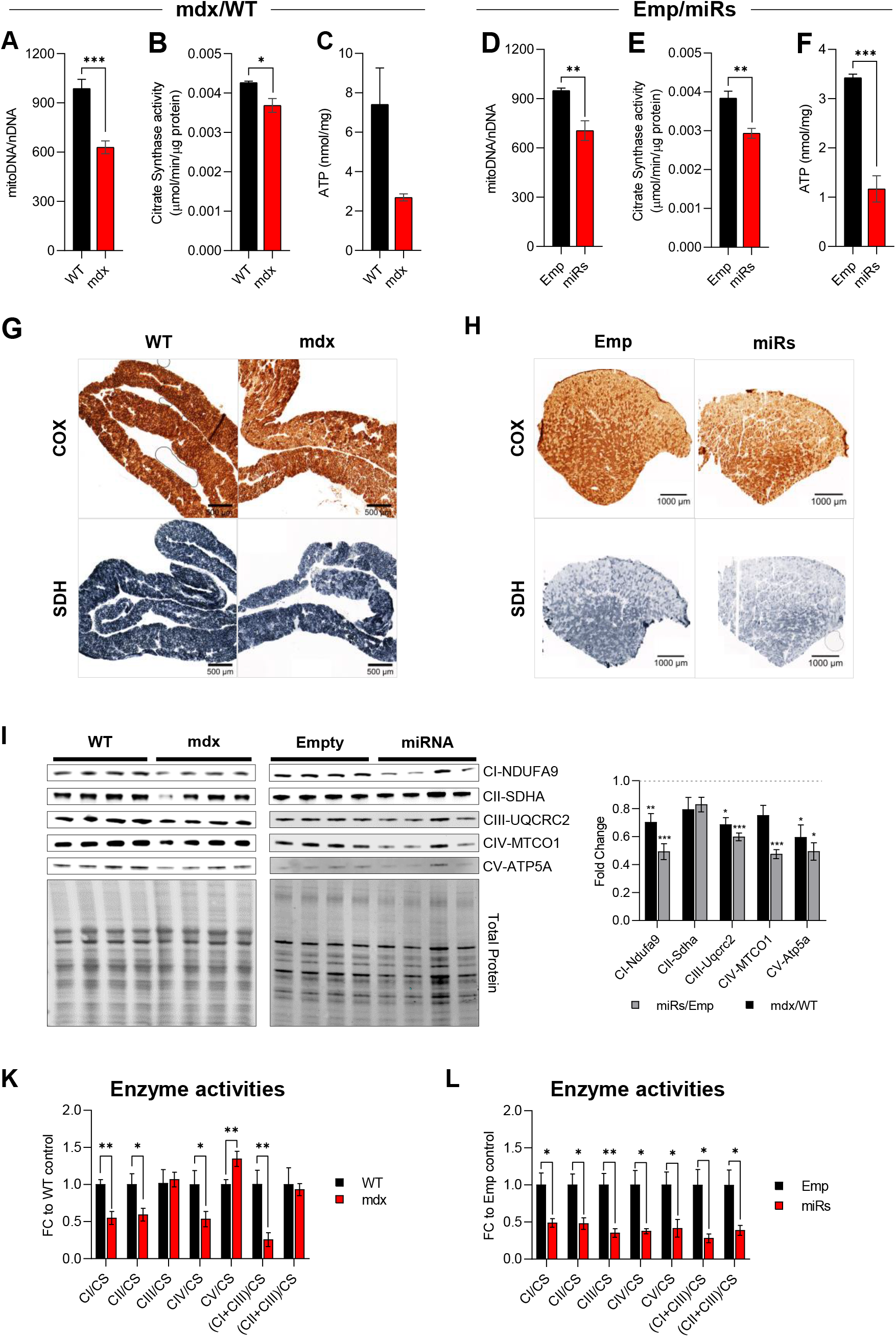
DD-miRNAs modulate mitochondrial metabolism in the dystrophic muscle. **A, D**. Levels of mitochondrial DNA in mdx versus WT diaphragm muscles (**A**, n=6) and miRs versus Emp TA muscles (**D**, n=4). **B, E**. Citrate Synthase activities in mdx versus WT diaphragm muscles (**B**, n=6) and miRs versus Emp TA muscles (**E**, n=6). **C, F**. ATP concentration in mdx versus WT diaphragm muscles (**C**, n=6) and miRs versus Emp TA muscles (**F**, n=6). **G**, **H**. Staining of COX (upper panels) and SDH (low panels) activities in mdx versus WT diaphragm muscles (**G**) and miRs versus Emp TA muscles (**H**). **I**. Western blots of five representative proteins for five OxPhos complexes comparing mdx versus WT diaphragm muscles and miRs versus Emp TA muscles (n=4), and their quantification (right). **K, L**. Activities of different OxPhos enzymes comparing mdx versus WT diaphragm muscles (**K**, n=5-6) and miRs versus Emp TA muscles (**L**, n=6). Data are presented as mean±SEM. Statistics were performed with Student t-test. *P < 0.05; **P < 0.01; ***P < 0.001.

A western blot analysis demonstrated reduced expression of proteins belonging to all four mitochondrial respiratory chain complexes and ATP synthase in the both mdx versus control diaphragms and DD-miRNAs versus control transduced TA muscle **(Figure 6I)**. Of interest, of the five proteins that were monitored in this experiment, two are potential targets of DD-miRNAs (SDHA is a potential target of miR-411-5p, and UQCRC2 is of miR-409-3p) while the other three (CI-NDUFA9, CIV-MTCO1, and CV-ATPA5) are not predicted to be direct targets of any DD-miRNA. It indicates that the downregulation of OxPhos proteins in the conditions of elevated DD-miRNA in the muscle involved a global downregulation of mitochondrial complexes, rather than (or in addition to) direct targeting by DD-miRNAs. We then tested the activities of the different mitochondrial complexes (normalized to Citrate Synthase activity (CS) – an indicator of mitochondrial mass) ^19^. Reduced complex activities were observed in complexes I, II, and IV in both systems, while complex III and V presented reduced activity only in the TA-DD-miRNAs muscles (**Figure 6K-L**). Significant reductions were also detected for the combined activity of complex I+III (NADH cytochrome c oxidoreductase) in both systems.

### Reduction in DD-miRNAs level increases OxPhos expressions and activities

Next, we wanted to ask whether mitochondrial activities could be enhanced by reducing DD-miRNAs expression. Since no viable mouse model exists with a silencing of the entire DD- miRNA cluster, we decided to create *in vitro* myogenic model of reduced DD-miRNAs expression. The M180 human induced pluripotent stem cell (hiPSC) line was selected, because it expresses high level DD-miRNAs throughout *in vitro* myogenesis ^20^. Since it was previously shown that maternal Dlk1-Dio3 expression, including DD-miRNAs, was abolished in IG-DMR−/− mouse embryonic stem cells ^21^ we deleted, in M180 iPS, the 11 kb IG-DMR by double-cut CRISPR/spCas9 strategy, with two sgRNAs target the two ends of IG-DMR (**Figure 7A**). Clonal selection was performed and biallelic deletion verified (Supplemental **Figure 5A**). IG-DMR^+/+^ (Ctrl) and IG-DMR^−/−^ (IG-KO) clones were differentiated *in vitro* into skeletal muscle myotubes (Supplemental **Figure 5B-C**). IG-KO clones showed normal myogenic differentiation capacity, by successfully converting into myotubes (Supplemental **Figure 5C**). However, while complete KO of DD-miRNA levels was seen in the iPS pluripotent state (D0), (Supplemental **Figure 5D**), only partial, but significant, reduction was observed in fully differentiated myotubes (Supplemental **Figure 5E**), suggesting that, while the DD-miRNA expression is controlled solely by IG initially, additional elements control their expression following maturation. We then investigated the mitochondrial responses in hiPSC-derived myotubes with reduced DD-miRNA levels. At the RNA level, IG-KO clones showed a global upregulation of transcripts of all five OxPhos complexes, although with variation between clones in the level of some genes (n=4, student t-test) (**Figure 7B**). Besides, increased mRNA levels in both IG-KO clones were also seen in two master regulators of mitochondrial biogenesis (Tfam and Ppargc1a), four DD-miRNAs predicted targets (**Figure 7B**), and two validated targets (Eif4g2, Usmg5) ^5^. Similarly, the protein levels of five representatives for five OxPhos complexes in the two IG-KO clones showed a global increase compared to the control (**Figure 8C**). More importantly, activities of all five OxPhos complexes in the KO clones increased significantly compared to control (**Figure 7F**). Accumulatively, these results support that reduced DD-miRNAs can enhance mitochondrial activity, particularly the OxPhos system in differentiated myotubes.

**Figure 7:**
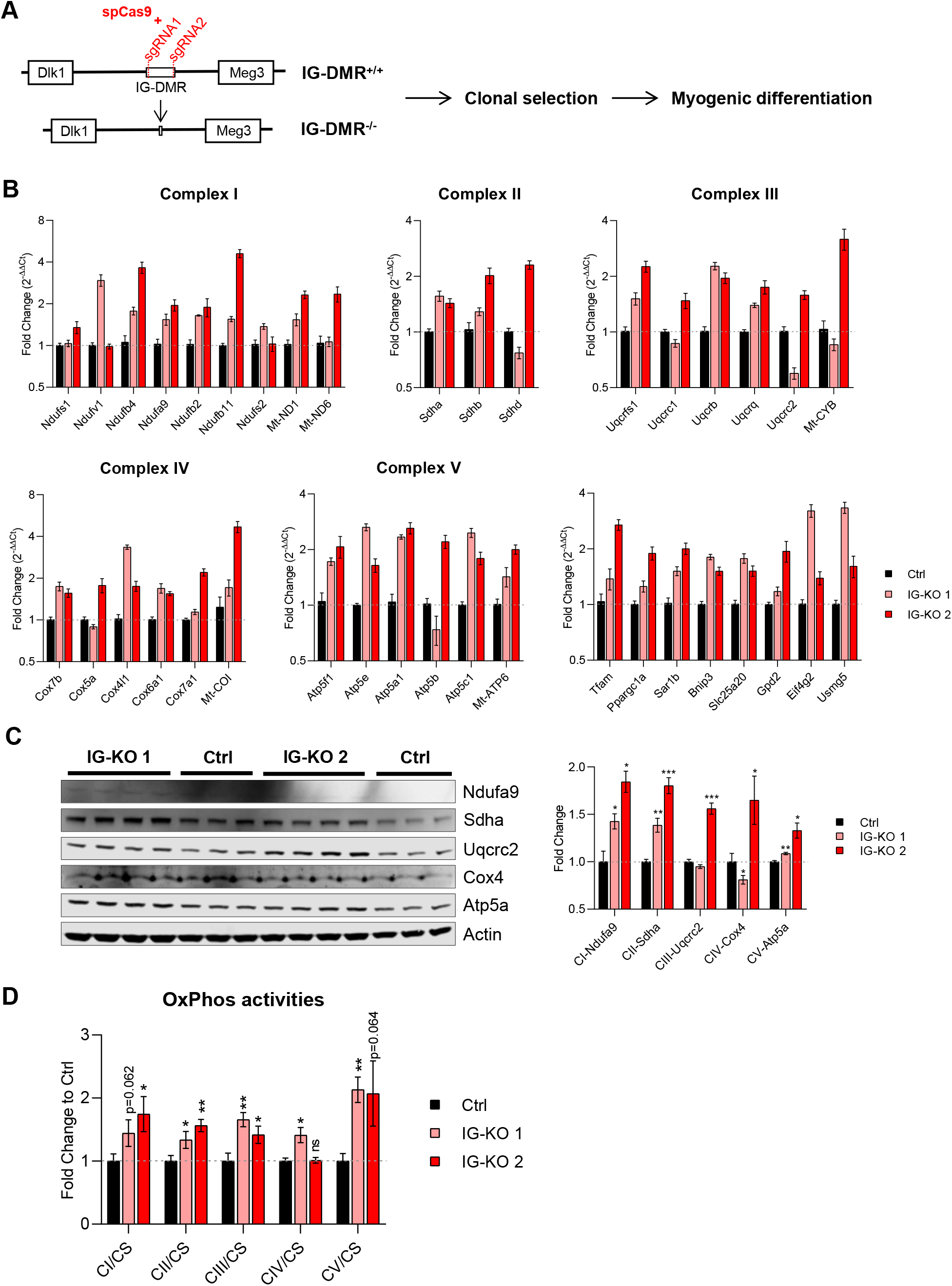
Knocking down DD-miRNAs results in increased OxPhos expression and activity. **A**. Diagram of IG-DMR deletion by CRISPR-Cas9. **B-D**. Analysis of DD-miRNAs knocked-down iPS clones compared to control (2 iPS clones, n=4). **(B)**, RNA level of selected transcripts of five OxPhos complexes and predicted DD-miRNA targets; **(C)**, Western blot of five representative proteins for five OxPhos complexes. **(D)**, CS-normalized activity of five OxPhos complexes. Data in **B-D** are presented as mean±SEM. Statistics were performed with Student t-test. *P < 0.05; **P < 0.01; ***P < 0.001.

## DISCUSSION

In previous investigations, we identified the upregulation of DD-miRNAs in the serum and the muscles of DMD patients and animal models ^3-5^. In the present report, we are bringing new evidences supporting a role for the DD miRNA cluster in the regulation of mitochondrial functions in DMD.

### Indications that circulating DD-miRNAs are produced and secreted by the regenerating muscle

In agreement with previous studies ^11,12^, we detected in adult mouse the highest DD-miRNAs expression in the brain, followed by the skeletal muscle. However, activation of expression in the mdx was detected only in the muscle. In addition, induced DD-miRNAs expression was observed in the serum of other mouse models for muscular dystrophy, without brain phenotype, making it unlikely that brain DD-miRNAs contributing significantly to the upregulation of circulating DD-miRNAs in muscular dystrophy. These data support strongly that regenerating myofibers are mostly responsible for the elevated levels of muscle and circulating DD-miRNAs in muscular dystrophy. The serum profiles of DD-miRNAs over different ages in GRMD dogs ^4^ and DMD patients ^3^ revealed a similar expression pattern to the mCK, of reduced dysregulation level beyond the age of 1 year in GRMD ^4^ and of 12 years in DMD ^3^. However, while the dropdown with age in DMD of serum mCK is thought to reflect reduced muscle mass and reduced myofiber degeneration, on the contrary, the dropdown with age of circulating DD-miRNAs is seem to best explained by reduced myofiber regeneration rate, which is consistent with the well-known exhaustion of the muscle regenerative capacity in DMD with evolution of the disease.

### Epigenetic control of DD-miRNA dysregulation in the dystrophic diaphragm

DNA methylation and post-translational modifications of core histones at differentially CpG methylated regions (DMRs) are the most characterized epigenetic mechanisms controlling Dlk1-Dio3 imprinting and expression ^6,21,22^. In non-muscular system, the increased expression of DD-miRNA was shown to be associated with DNA hypomethylation ^23,24^, and aberrant histone modifications ^25^at these DMRs. Epigenetic control was shown to play a key role in the regulation of muscle functions in muscular dystrophy ^26,27^. We thus hypothesized that in DMD, epigenetic control may mediate the dysregulation of DD-miRNAs. To test the hypothesis that, in the muscular system, DD-miRNAs might be subjected to epigenetic control, we took advantage of the C2C12 myoblasts. Global epigenetic analysis was performed with the non-specific inhibitors of DNA methylation, (5-Azacytidine), and histone deacetylation (Givinostat and TSA) and confirmed this possibility. Gene expression analysis in the mdx diaphragm validated global changes in genes, which are involved in an epigenetic control. The analysis of epigenetic marks at IG-DMR and Meg3-DMR supported that the decreased expression of the repressive H3K27me3 marker and the decreased DNA methylation level (of IG-DMR only) are the direct causes of DD-miRNAs upregulation in the dystrophic diaphragm.

### The DD-miRNA cluster regulates mitochondrial OxPhos in the dystrophic muscle

Rather than a hypothesis-driven approach, we undertook here a strategy of a non-supervised investigation, consisting of the *in vivo* co-overexpression of a 14 DD-miRNAs, and global transcriptomic bioinformatics analysis. This analysis pointed toward lipid metabolism and mitochondrial functions as the most dysregulated pathways downstream to the DD-miRNAs, supporting experimentally strongly that the mitochondrial OxPhos system is indeed the most affected pathway in the dystrophic muscle by the cooperative activity of the DD-miRNAs. Since the activation of DD-miRNAs was also observed in the muscle and serum of three LGMD mouse models and during regeneration of normal muscle after Notexin-induced injury, the involvement of DD-miRNAs in mitochondrial functions might be associated with diverse situations of muscle injury and regeneration.

### *In vivo* validation of control of mitochondrial OxPhos by DD-miRNAs

We wondered to what degree the dystrophic changes in the mdx muscle might be explained by the upregulation of DD-miRNAs. To answer this question, we overexpressed 14 DD-miRNAs *in vivo* in the muscle and compared transcriptomic, protein level, and mitochondrial activity of the mdx muscle to the DD-miRNA overexpressing C57Bl/6 muscle. We detected a high overlap between omics changes in the normal muscle overexpressing DD-miRNAs, and in the dystrophic muscle, naturally having elevated DD-miRNA levels. Of particular interest, the mitochondrial OxPhos, which is the highest bioinformatics-predicted target pathway for DD-miRNAs, was experimentally detected as the most downregulated pathway in both conditions. This observation strongly supports that the DD-miRNAs are direct repressive mediators of OxPhos system in the dystrophic muscle. The upregulated DD-miRNAs are expected to downregulate target genes in the dystrophic muscle. As expected, therefore, we found a high-level overlap in the downregulated transcripts (which are the direct targets of DD-miRNAs), in the normal muscle overexpressing DD-miRNAs, and in the dystrophic muscle. Surprisingly, a high overlap was found also in the upregulated transcripts, supporting that non-direct effects of DD-miRNAs may similarly occurs in the two systems. Taken together, the data support that DD-miRNAs upregulation plays important role in mediating omics changes and mitochondrial activity in the dystrophic muscle. The direct evaluation of the silencing of all DD-miRNAs in the mdx mouse was not possible, since the knocking out of the entire DD-miRNA cluster is embryonically lethal. We selected therefore an *in vitro* iPS-based strategy for the modeling of DD-miRNAs silencing in the muscle. The CRISPR-Cas9 induced deletion of the IG-DMR, led to a drastically reduced DD-miRNA expression in the *in vitro* differentiated skeletal myotubes derived from a human iPS cell line. The analysis of mitochondrial gene expression and activity in myotubes derived from two independent clones confirmed that DD-miRNA might control mitochondrial OxPhos in the muscular system. Thus, the present study linked DD-miRNAs with the control of mitochondrial activity in the regenerating myofiber.

DD-miRNAs are highly expressed in the quiescence muscle stem cell and down-regulated upon its activation ^13^ or in dystrophic satellite cells (**Figure 1E**), however the biological function of DD-miRNAs in muscle stem cells is yet unknown. Of note, it has recently been suggested that activation of the satellite cells in the exercised muscle is OxPhos–dependent ^28^. It is tempting to speculate that by OxPhos inhibition DD-miRNAs are involved in the maintenance of stem cells quiescence in the resting muscle, whiles DD-miRNAs downregulation is required for the activation of stem cells in the exercising or regenerating muscle.

### Updated model for mitochondrial dysfunction in DMD

It has been suggested long ago, that the entry of Ca^2+^, which is provoked in muscular dystrophy by sarcolemma instability, is followed by a pathological cascade that includes Ca^2+^ impact on mitochondria, leading to myogenic degeneration^29^. This proposition for mitochondrial dysfunction in muscular dystrophy is still widely accepted today ^30,31^. We have recently demonstrated that increased miR-379 expression in the dystrophic muscle interferes with the OxPhos system and ATP production. Based on this discovery, we proposed a modified model for mitochondrial dysfunction in DMD. Our updated model for mitochondrial participation in the dystrophic cascade supports that in muscular dystrophy, not only miR-379, but also many other DD-miRNAs are coordinately upregulated and cooperatively affect mitochondrial functions in the dystrophic muscle.

In summary, the present investigation exposes a fundamental mechanism of mitochondrial adaptation in the dystrophic muscle and therefore opens a new perspective for therapeutic intervention.

## Author contributions

**AVH** designed and performed experiments, analyzed results and wrote the paper. **NB, PS, JP, KC, EG, EM,** performed experiments and analyzed results. **MS** designed experiments and analyzed results. **IR** and **DI** designed experiments, analyzed results, managed the project and wrote the paper.

## Acknowledgment

The authors are Genopole’s members, first French biocluster dedicated to genetic, biotechnologies and biotherapies. We are grateful to the “Imaging and Cytometry Core Facility” and to the *in vivo* evaluation, services of Genethon for technical support, to Ile-de-France Region, to Conseil Départemental de l’Essonne (ASTRE), INSERM and GIP Genopole, Evry for the purchase of the equipment. We are grateful to the Myobank, the tissue bank of the Association Française contre les Myopathies (AFM), and the Myology Institute (Paris, France) for providing DMD and healthy control skeletal muscle biopsies. This study was financially supported by the AFM; by the Institut National de la Sante et de la Recherche Medicale (INSERM); the University Pierre et Marie Curie Paris 06; the Centre National de la Recherche Scientifique (CNRS).

## Supplemental material

**Supplemental Figure 1:**
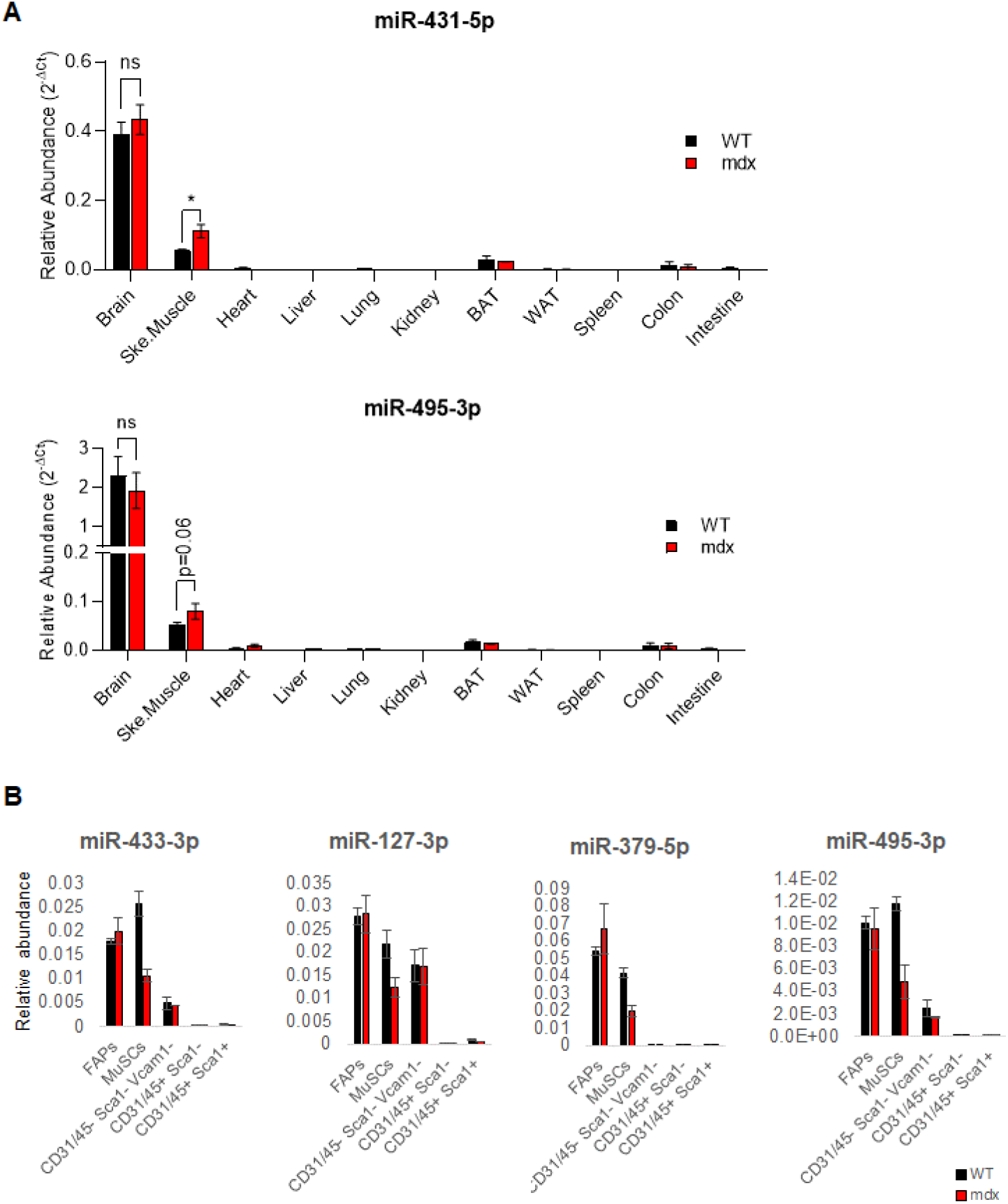
DD-miRNAs expression level in different organs, tissues and skeletal muscle cellular subpopulations. **A.** Comparison of expression levels of 2 representatives DD-miRNAs (miR-431-5p and miR-495-3p) between different organs from 5-week-old wild-type and mdx mice (n=3-4). **B**. Comparison of expression levels of 4 representative DD-miRNAs between different FACS-sorted cell populations from 5-week-old mdx mice and its healthy controls (n=3). Muscle mono-nucleated cells (MMNC) fractions were FACS-sorted by using CD31, CD45, Sca1 and Vcam1 markers from hind limb muscles of mdx and control mice. We quantified the expression of four representative DD-miRNAs (n=3) in five subpopulations that are typically found in skeletal muscle, which are the hematopoietic cells (CD45+), endothelial cells (CD31+), fibro-adipocyte progenitors (FAPs) (CD31-/CD45-/Sca1+), satellite cells (CD31-/CD45-/Sca1-/Vcam1+), and cells that are negative for all four markers (CD45-/CD31-/Sca1-/Vcam1-). Expression of DD-miRNAs was detected in FAPs, satellite cells, and CD31/CD45/Sca1/Vcam1 negative cellular fraction. Comparison between the mdx and WT cells showed that DD-miRNAs were expressed to similar levels in the FAPs and in the negative cell population. In contrast, a lower DD-miRNAs expression was detected in satellite cells derived from the mdx mouse compared to control, which is consistent with the activated state of satellite cells of the mdx mouse, and with the fact that DD-miRNAs are known to be downregulated in activated satellite cells ^1^. Data are presented as mean±SEM. *P < 0.05; **P < 0.01; ***P < 0.001.

**Supplemental Figure 2:**
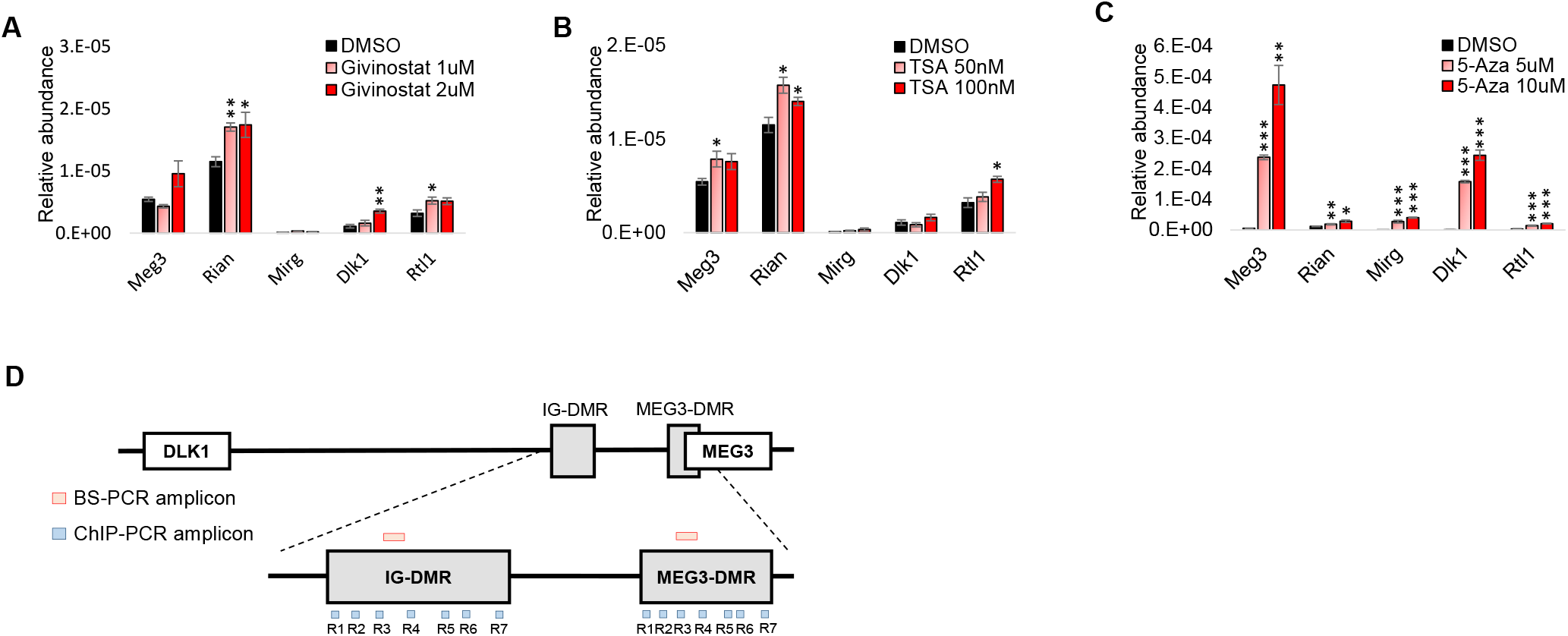
Epigenetic drugs affect the expression of Dlk1-Dio3 transcripts in myogenic cells. Relative expression of maternal and paternal Dlk1-Dio3 transcripts in C2C12 cells treated by Givinostat (**A**), Trichostatin A (**B**), and 5-azacytidine (**C**) compared to DMSO control (n=4). **D.** Illustration of the PCR amplicons in Figure 2A-B in the context of the DLK1-Dio3 locus. TSA, Trichostatin A; 5-Aza, 5-azacytidine; BS-PCR, PCR from bisulfite-converted DNA; ChIP, Chromatin Immuno-Precipitation. Data are presented as mean±SEM. *P < 0.05; **P < 0.01; ***P < 0.001.

**Supplemental Figure 3:**
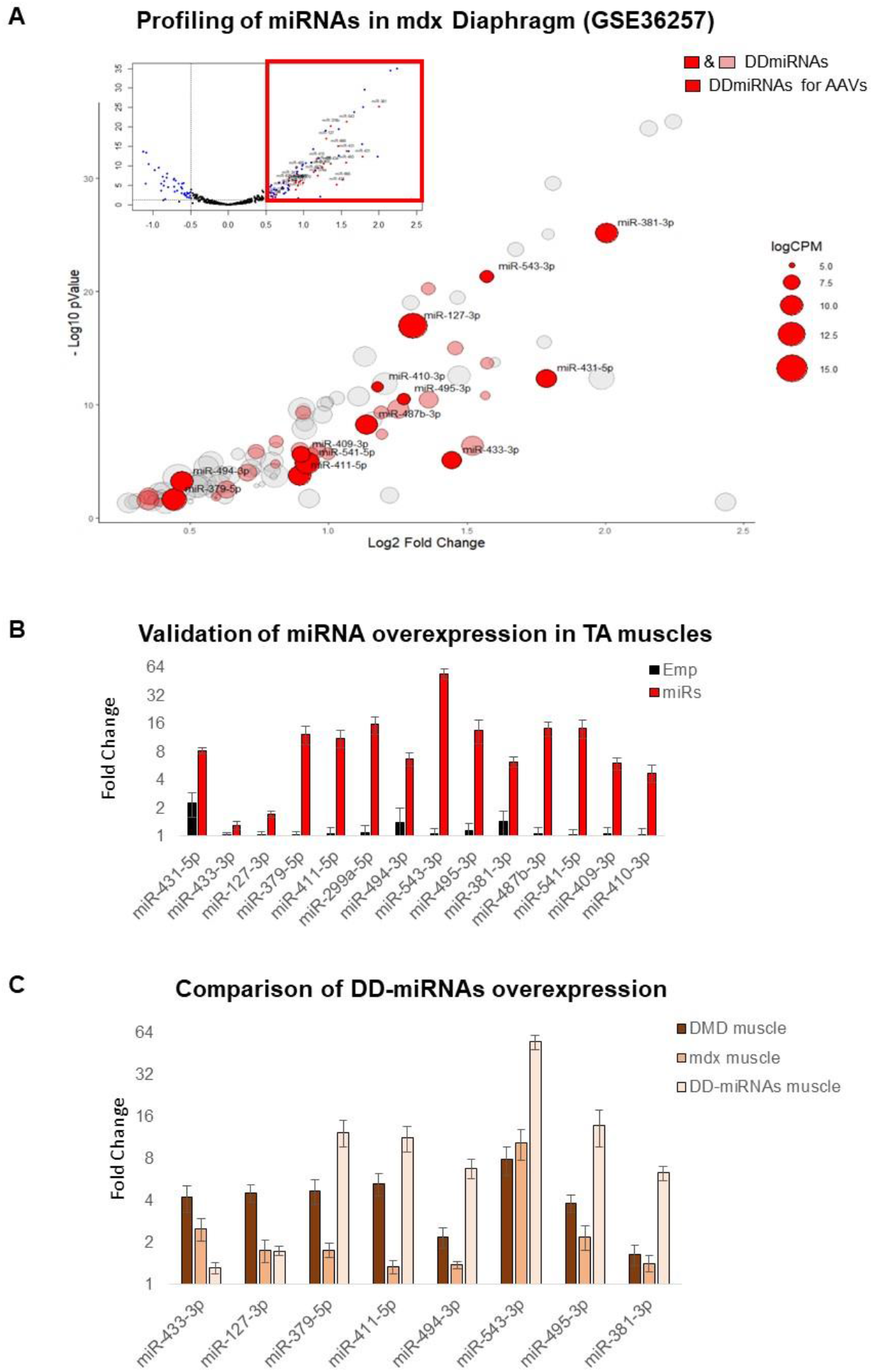
Selection and *in vivo* overexpression of 14 DD-miRNAs in skeletal muscle. **A.** Volcano plot of miRNA profiling of the diaphragm of 8-week-old mdx and control mice (upper part), and a zoom-in of significantly up-regulated miRNAs in mdx muscle (lower part) (data was taken and reanalyzed from GEO: GSE36257 ^2^. Expression levels are presented by the size of the circle. DD-miRNAs are colored (pink or red). MiRNAs in red were selected for the overexpression by AAV vectors in the present study. **B**. Validation of DD-miRNAs overexpression in the treated mice. TA muscles were analyzed one month after injection with AAV-DD-miRNAs (miRs) compared to AAV-Empty control (Emp) (n=6). **C**. Comparison of DD-miRNAs overexpression in different systems: muscle biopsies from DMD patients compared to healthy controls (Figure 1A), diaphragm muscle of mdx mice compared to C57Bl10 control (Figure 1B), and DD-miRNAs overexpression in the TA muscle of a healthy mouse by AAV vectors. Data in **B-C** are presented as mean±SEM.

**Supplemental Figure 4:**
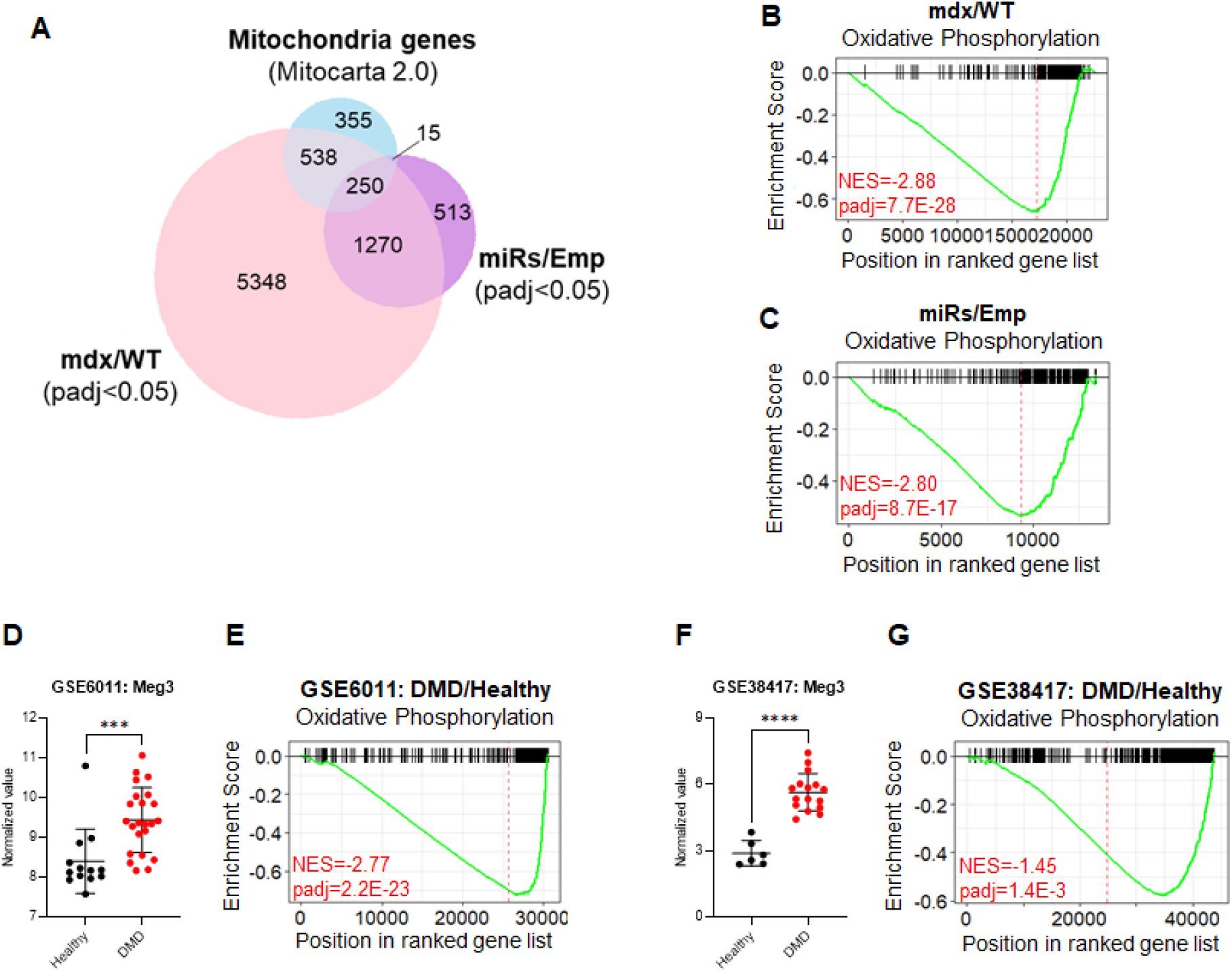
DD-miRNAs affect mitochondrial functions in the dystrophic muscle. **A.** Venn diagram of mitochondria-related genes (Mitocarta 2.0) and of significantly dysregulated genes, in mdx/WT and miRs/Emp transcriptomes. **B-C**. GSEA analysis of OxPhos gene set comparing mdx versus WT diaphragms (**A**) and miRs versus Emp TA muscles (**B**). **D-G**. Expression level of Meg3 transcript and GSEA analysis of OxPhos gene set in muscle biopsies of DMD patients compared to healthy controls in two different datasets. Data was taken and reanalyzed from GEO GSE6011 ^3^ (**D-E**) and GEO GSE38417 (**F-G**). Data are presented as mean±SD. *P < 0.05; **P < 0.01; ***P < 0.001.

**Supplemental Figure 5:**
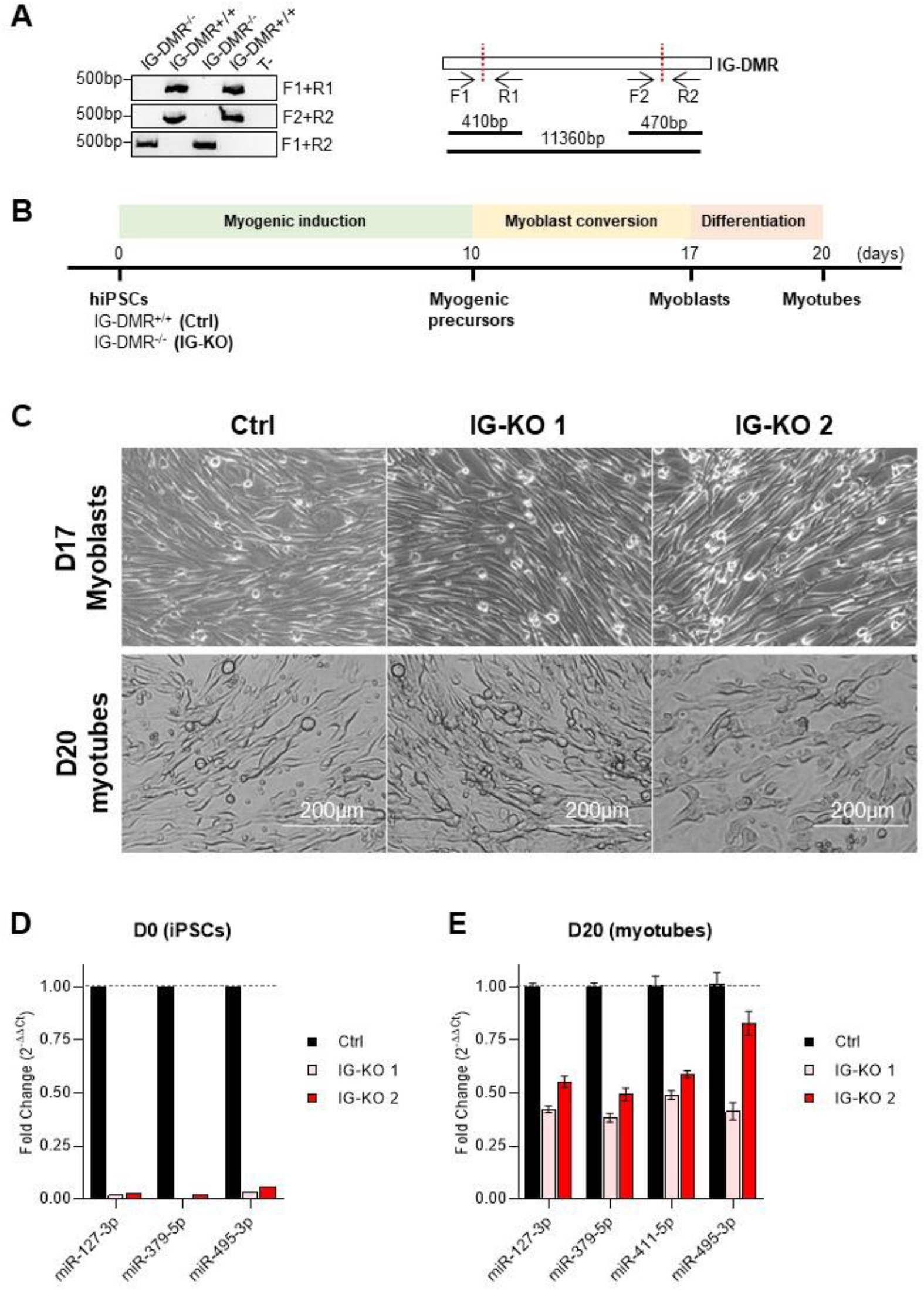
Biallelic deletion of IG-DMR in hiPSCs. **A.** Confirmation of 11-kb IG-DMR deletion in different IG-DMR^−/−^ clones by PCR using different sets of primers (left). Positions of primers used in PCR are illustrated (right). **B**. Diagram of myogenic differentiation methods for hiPSCs. **C**. Representative images of D17 myoblasts (upper panels) and D20 myotubes (lower panels) of different IG-DMR^−/−^ and control clones. **D-E**. Relative expression levels of representative DD-miRNAs at D0 (**D** - hiPSCs) and D20 (**E** - myotubes) in 2 IG-DMR^−/−^ clones compared to control (n=4). Data are presented as mean±SEM.

## Supplemental Tables

**Supplemental Table 1:**
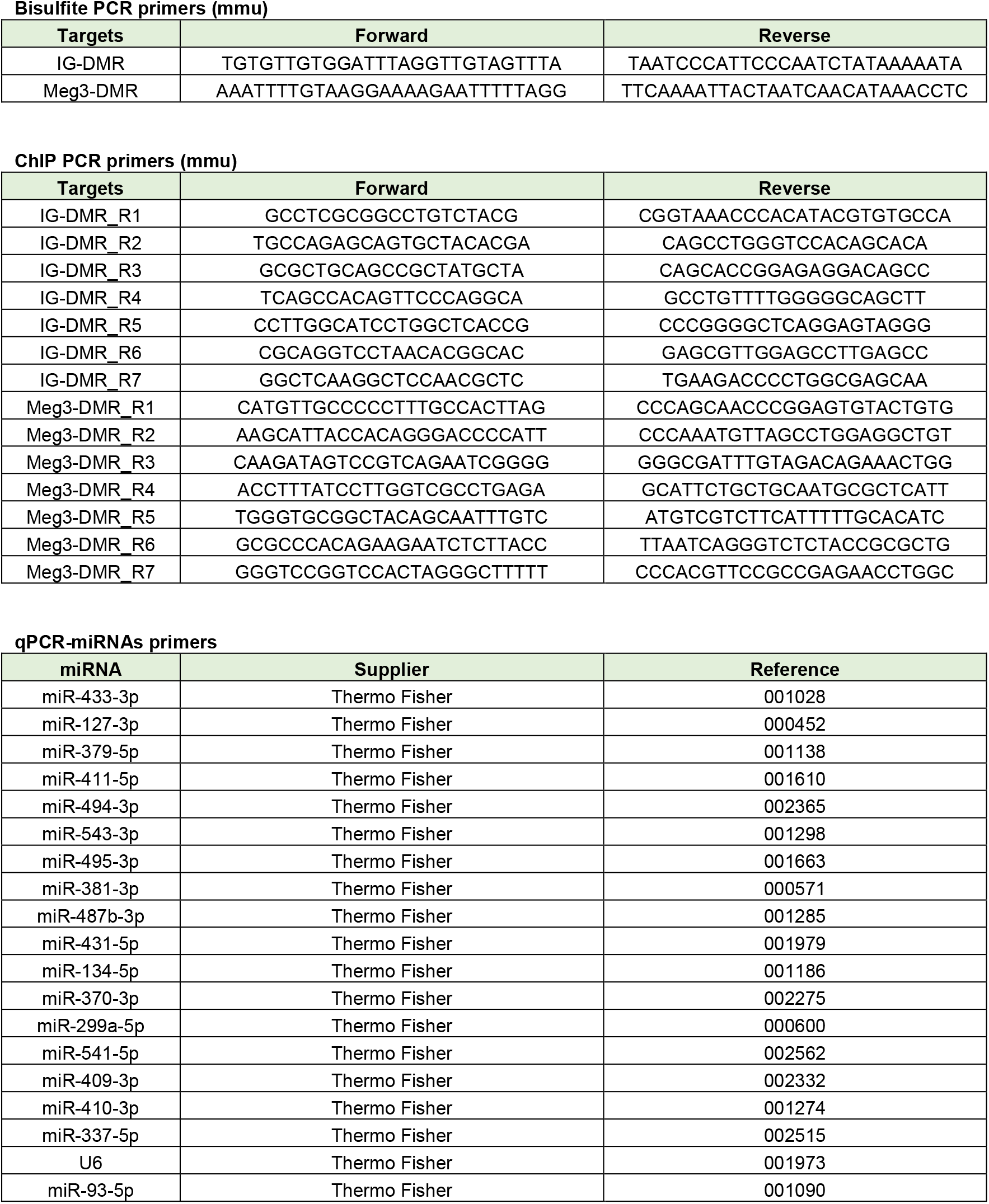
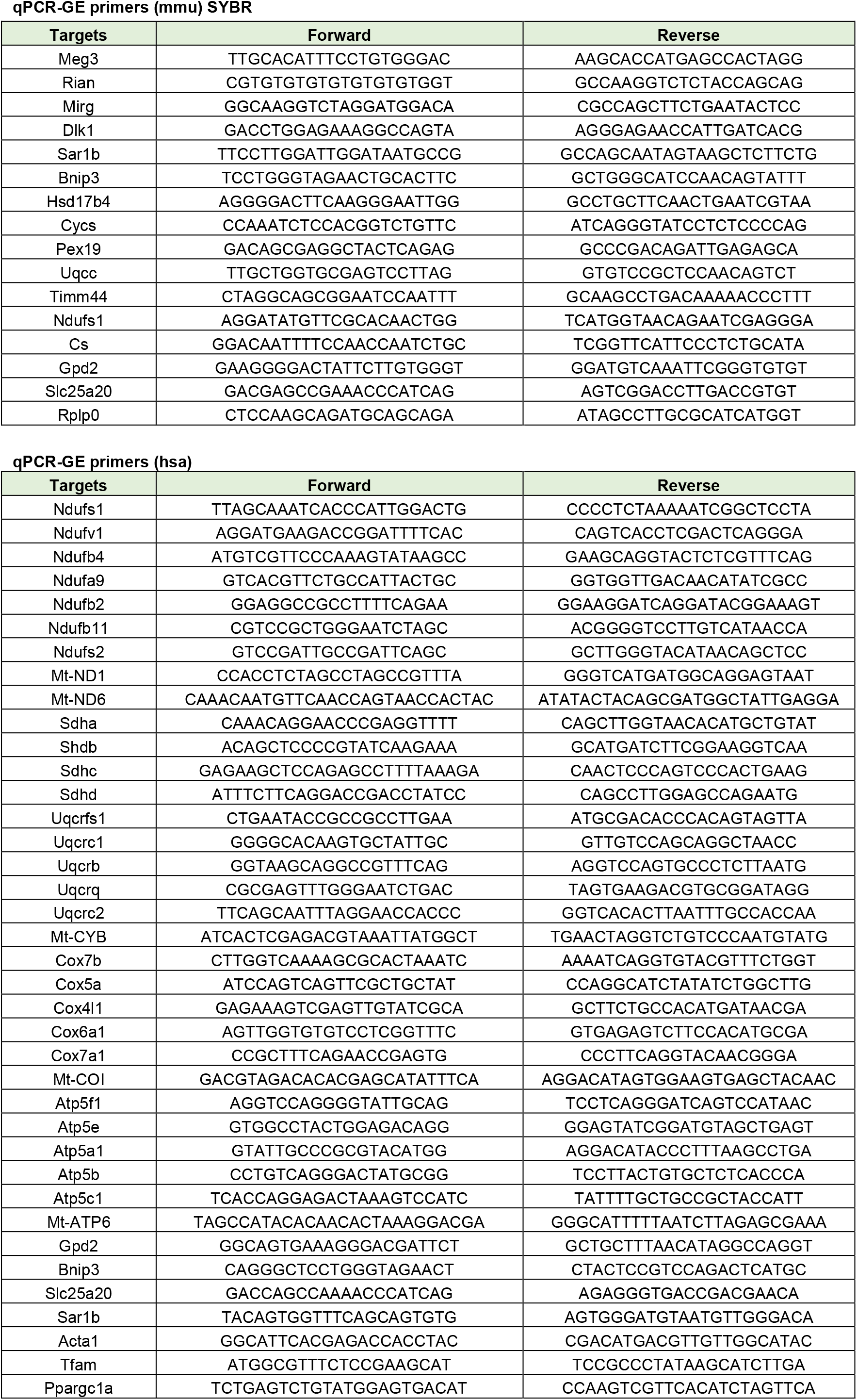
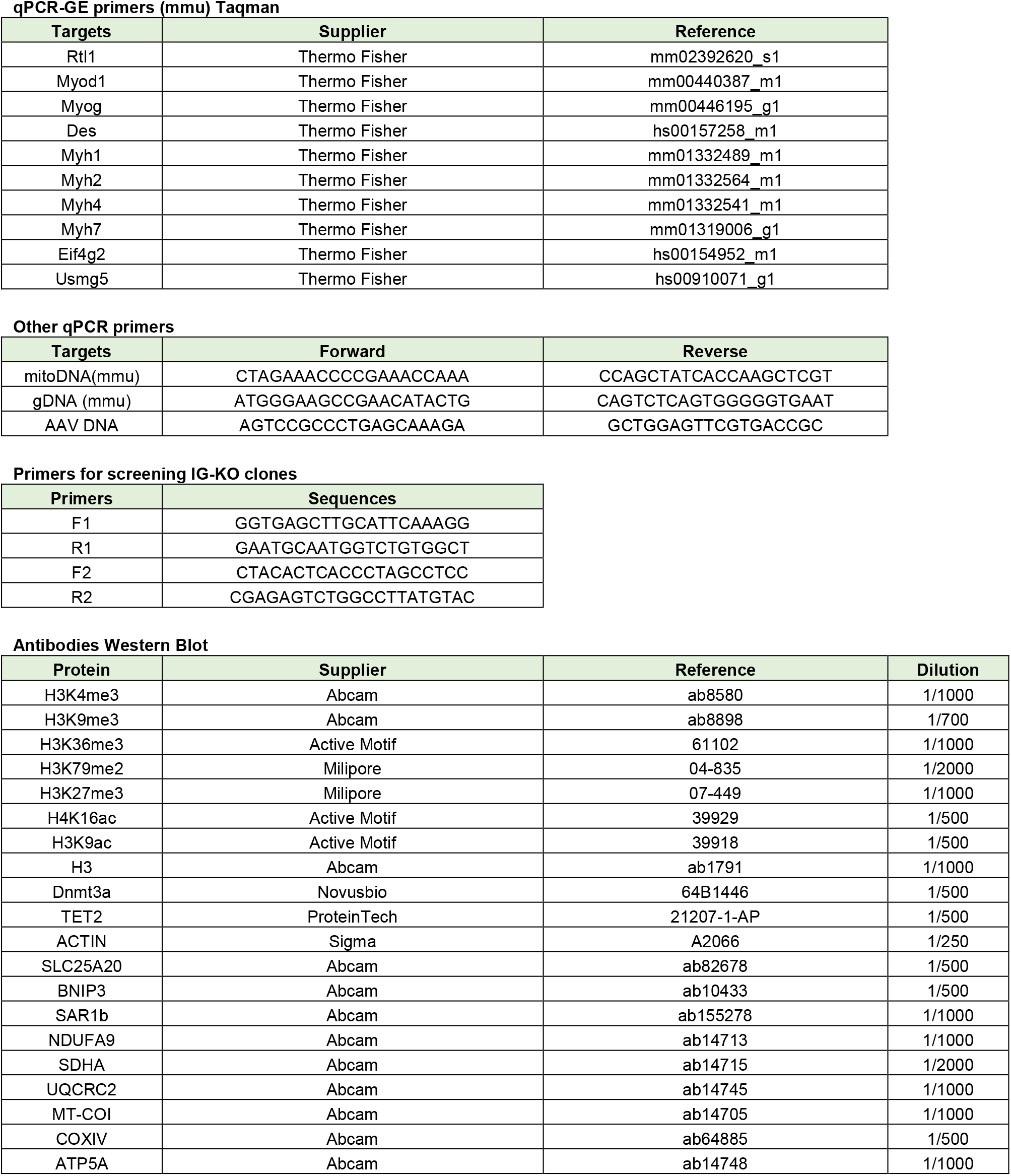
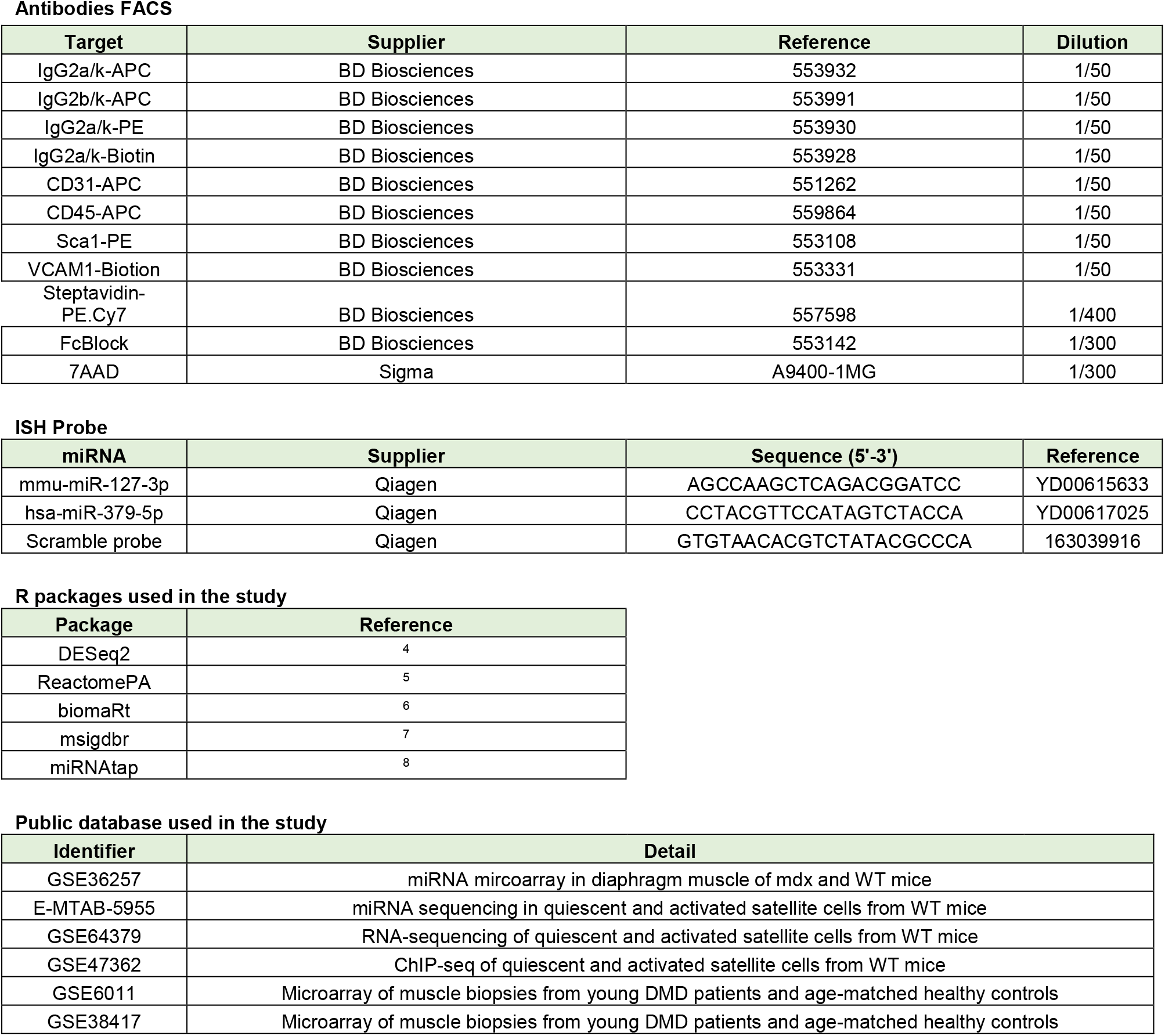
List of primers, antibodies, reagents, R packages, and public databases used in the present study.

**Supplemental Table 2:**
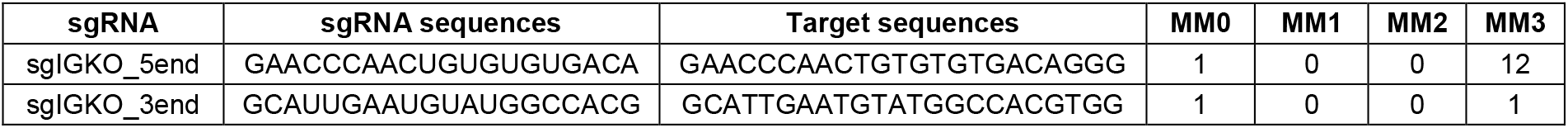
List of sgRNAs used in the present study. Selection of sgRNAs were done by GPP sgRNA Designer ^9^. Number of off-targets (MM0-3) were further identified by CHOPCHOP ^10^. MM0, −1, −2, −3: number of off-targets with 0, 1, 2, 3 mismatches, respectively.

**Supplemental Table 3:**
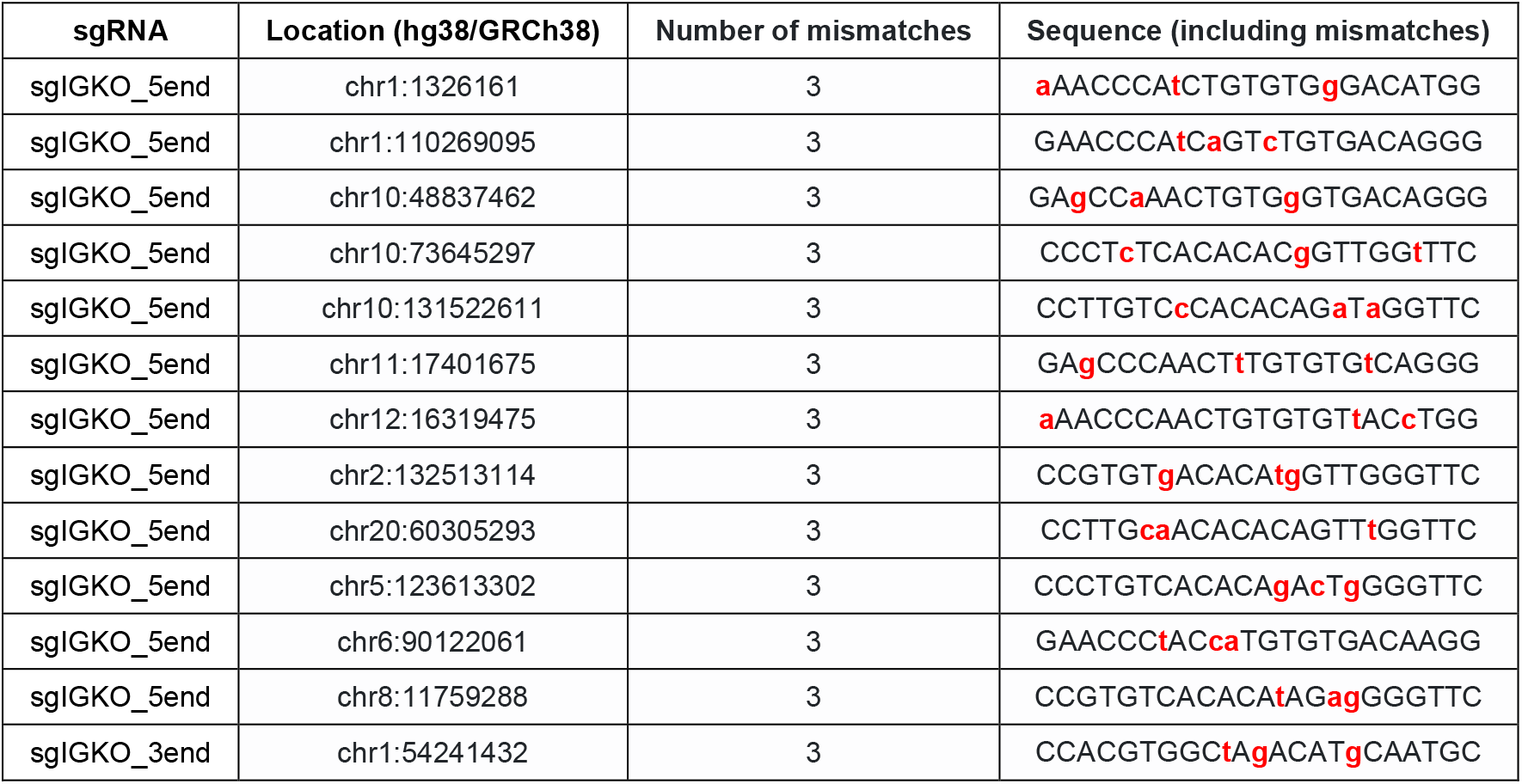
List of off-targets associated with sgRNAs used in this study. Off-targets were identified by CHOPCHOP ^10^. Mismatches of off-target sequences are colored in red.

**Supplemental Table 4:** Selection of DD-miRNAs for overexpression in muscle. Fourteen miRNAs were selected to be overexpressed in the TA muscle of the WT mouse. The choice of miRNAs was based on their expression and dysregulation levels in dystrophic serum, skeletal muscle and quiescent satellite cells. Overexpression data is provided in the excel file **“Supplemental Table 4”**

**Supplemental Table 5:** Gene Set Enrichment Analysis of transcriptomes from dystrophic muscle and DD-miRNAs overexpressed muscles. The data is provided in the Excel file **“Supplemental Table 5”**

**Supplemental Table 6:** List of bioinformatics predicted target genes for the selected 14 DD-miRNAs. Five prediction tools (DIANA, Targetscan, PicTar, Miranda, and miRDB) were used, generating 5295 candidate targets predicted by at least two different tools. This list of 5295 candidates was crossed with experimental data of transcripts that are downregulated in the injected TA muscle and in the mdx diaphragm. The analysis resulted in 269 DD-miRNAs predicted targets, which is provided in the Excel file **“Supplemental Table 6”**

### Extended materials and methods

#### Muscle biopsies

Human skeletal muscle tissues were obtained from Myobank, the tissue bank of the Association Francaise contre les Myopathies (AFM; https://www.institut-myologie.org/en/recherche-2/myobank-afm/). Open skeletal muscle biopsies were performed after informed consent, according to the Declaration of Helsinki. Muscle biopsies included in this study were derived from paravertebral (two controls and two DMD), gluteus, and tensor fasciae latae (one each control and one DMD) muscles. Details of patients and the biopsies are presented in **Table 1**.

**Table 1:**
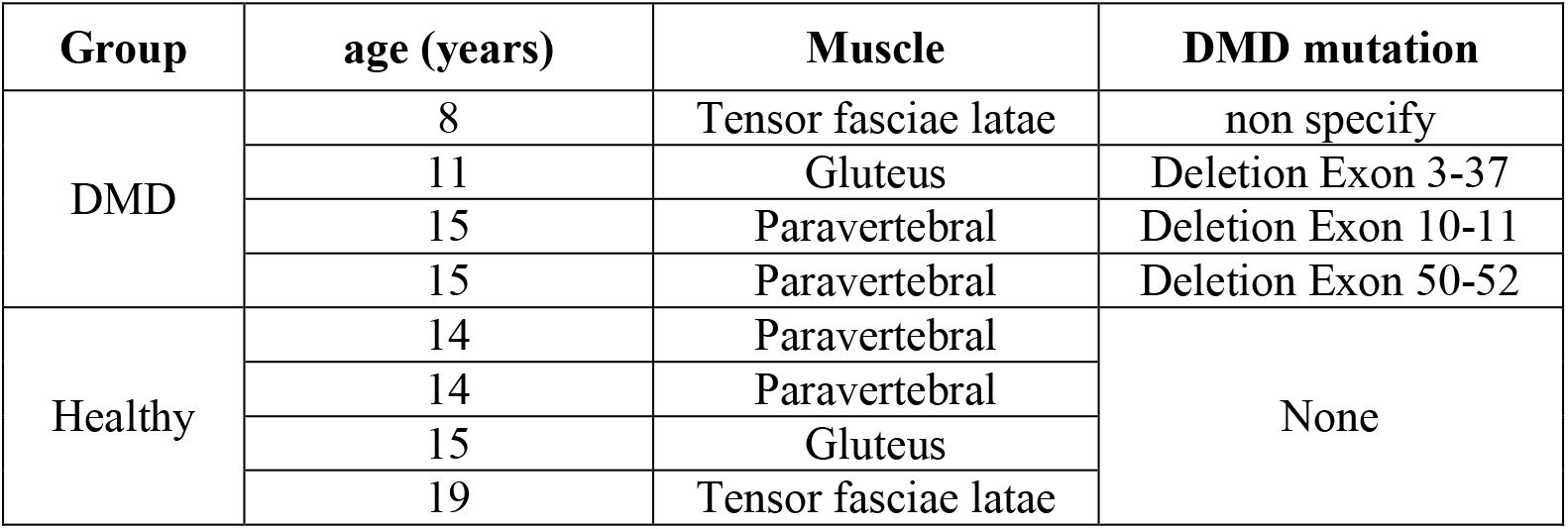
Human muscle biopsies used in the present study.

#### AAV construction and production

The 14 pre-miRNA sequences were obtained from UCSC website (https://genome-euro.ucsc.edu), spanning 100 nucleotides before and after the mature miRNA sequences. The selected pre-miRNA sequences were then arranged consecutively which respect the genomic sequences of the miRNAs. Two or three pre-miRNAs were used per AAV construct. The constructed sequences were then synthesized (Genewiz) and subcloned into a donor plasmid (AAV-CMV-eGFP) by classical molecular biology technique. AAV9 viral vectors production was performed as already described ^11^. Titration was performed by qRT-PCR using primers corresponding to the sequences of AAV ITR or eGFP. Sequences of PCR primers can be found in **Supplemental Table 1**.

#### Satellite cell isolation from skeletal muscles and their *in vitro* differentiation

Satellite cells were isolated from limb muscles as described ^12^, with some small modifications. All hind limb muscles of 5-weeks old mice were collected and minced by using scissors. The samples were then put in the digestion medium of DMEM with 3U/ml Dispase II (Thermo Fisher Scientific), 0.5 U/ml Collagenase A (Sigma), 0.2% BSA (Sigma), and 1X Pen-Strep (Thermo Fisher Scientific) for two hours at 37°C with gentle shaking. The lysates were then passed through successive strainers (100, 70, and 40 µm) to eliminate fibers and debris. Subsequently, red blood cells were removed from the samples by Versalyse (Beckman Coulter, Reference A09777). The mono-nucleated cells were then blocked by Mouse BD Fc Block (BD Pharmingen, Reference 553142), stained with CD31-APC, CD45-APC, Sca1-PE, Vcam1-PE.Cy7 and viability marker 7AAD. The corresponding isotype controls were used in parallel to determine the non-specific binding of antibodies. Details of antibody panel used in FACS sorting are presented in **Supplemental Table 1**. The cells were then FACS-sorted by Astrios Cell Sorter (Beckman Coulter). Satellite cells, marked as CD31^−^ CD45^−^ Sca1^−^ Vcam1^+^, were directly placed in the proliferating medium containing Ham’s F-10 (Hyclone) supplemented with 10% horse serum and 2.5 ng/ml bFGF (PeproTech, Reference 100-18B), and incubated at 37°C, 5% CO2 for 3 days. Proliferated cells were then switched into a differentiation medium containing DMEM and 5% Horse Serum (Gibco) and kept at 37°C, 5% CO2 for 5 days.

#### Bisulfite Conversion and DNA methylation analysis

Genomic DNA of muscle tissues was extracted by Phenol:Chloroform:Isoamyl Alcohol standard protocol. Purified DNA was then treated with sodium bisulfite by using EpiTech Bisulfite Kit (Qiagen, Reference 59104) according to the manufacturer’s protocol. Subsequently, bisulfite converted DNA was amplified by PCR using OneTaq HotStart DNA Polymerase (New England Biolabs, Reference M0481) with touch-down thermocycler program. Each PCR reaction was performed in triplicate. The amplified PCR products were subsequently subjected to Sanger sequencing (Genewiz), and the levels of modification at single nucleotides were quantified by EditR ^13^. The primers used in bisulfite PCR are detailed in **Supplemental Table 1.**

#### Chromatin immunoprecipitation

Chromatin immunoprecipitation (ChIP) of frozen muscle biopsies was performed using MAGnify Chromatin Immunoprecipitation System (Invitrogen) according to the manufacturer’s protocol with minor modifications. First, 50-100 mg muscle tissues were minced quickly on ice by scissor, and homogenized thoroughly by sequential uses of 16G, 18G, and 21G needle. Then, homogenates were immediately fixed in 1% formaldehyde for 10 minutes, quenched in 0.125M glycine, and washed twice in ice-cold PBS. Subsequently, the lysates were sonicated for 15 cycles of 30 seconds ON, 30 seconds OFF using Bioruptor Pico (Diagenode). Diluted lysates at this step were used as Input Control. The magnetic beads, which were already incubated with antibodies, were then incubated with the sheared chromatin for 2 hours at 4°C on the orbital shaker. After that, the formaldehyde crosslinks of the bound chromatin and Input Control were reversed and un-crosslinked DNA could be purified. Purified DNA was subjected to qPCR using SYBR Green PCR assays (Thermo Fisher Scientific). Each PCR reaction was performed in duplicate. Level of Input Control DNA was used to normalize the data across samples. Besides, ChIP reactions using rabbit and mouse IgG were also used as negative control. The antibodies and PCR primers used in ChIP-PCR were detailed in **Supplemental Table 1**. The location of ChIP-PCR primers is shown in **Supplemental Figure 2E**.

#### RNA expression analysis

Total RNA was extracted from frozen muscles or cells using Trizol (Thermo Fisher Scientific). DNA contamination from RNA samples was subsequently removed by TURBO DNA-free kit (Thermo Fisher Scientific). For the measurement of *miRNA expression*, 500 ng of total RNA were reverse-transcribed using TaqMan™ MicroRNA Reverse Transcription Kit (Thermo Fisher Scientific). Quantitative PCR was performed using LightCycler 480 system (Roche), with the Applied Biosystems TaqMan MicroRNA Assays (Thermo Fisher Scientific), according to the manufacturer’s protocol. Each PCR reaction was performed in duplicate. Results obtained with miR-93-5p or U6 small nuclear RNA were used to normalize data across samples. For the measurement of *gene expression*, 1000 ng of total RNA was reverse-transcribed using a mixture of random oligonucleotides and oligo-dT and RevertAid H Minus First Strand cDNA Synthesis Kit (Thermo Fisher Scientific). Quantitative PCR was performed using LightCycler 480 system (Roche) with the SYBR™ Green PCR assays (Thermo Fisher Scientific) or Taqman Gene Expression Assay (Thermo Fisher Scientific). Each PCR reaction was performed in duplicate. Results obtained with Rplp0 (for muscle samples) or Acta1 (for *in vitro* cells) were used to normalize the data across samples. Relative expression fold change was calculated using 2^−ΔΔCT^ method, as previously described ^14^. Primers used in the study are presented in **Supplemental Table 1.**

#### RNA sequencing and transcriptomic analysis

RNA quality was first examined by Bioanalyzer 2100 (Agilent). Samples with RNA integrity number greater than 7.5 were then subjected to RNA sequencing (Genewiz). The sequencing libraries were prepared using the Stranded Total RNA Library Prep Kit (Illumina) and sequenced according to the Illumina protocol by NovaSeq instrument (Illumina), resulting in approximately 20M paired-end reads per library. Filtration and quality control was done by fastp ^15^. The pair-end reads were subsequently mapped into GRCm38/mm10 genome by HISAT2 ^16^, and count tables were produced by FeatureCount ^17^. Differentially expressed genes were identified by DESeq2 R package ^4^. Pathway analysis was performed in R-Studio (version 4.0.3), either by over-representation methods using Gene Ontology and ReactomePA or functional class scoring using Geneset Enrichment Analysis (GSEA). The list of R packages used in the analysis can be found in **Supplemental Table 1**.

#### Cell culture and *in vitro* study

Myoblast C2C12 cells were maintained in the proliferating medium containing DMEM (Thermo Fisher Scientific) supplied with 10% Fetal Bovine Serum and 1X Gentamycin at 37°C, 5% CO_2_. Human induced pluripotent stem cells (hiPSCs) were maintained in mTeSR medium (Stemcell Technologies, Reference 05850) on Matrigel-coated plate (Corning, Reference 356230). To differentiate hiPSCs into skeletal muscle linage, 2D directed differentiation protocol was used as previously described ^18^. The hiPSCs were placed in three consecutive defined media as followed: SKM01 (AMSbio) from day 0 to 10, SKM02 (AMSbio) from day 10 to 17, and SKM03 (AMSbio) from day 17 to 20 (**Supplemental Figure 5B**).

#### Genomic deletion using the CRISPR/Cas9 and screening for bi-allelic deletion clones

Single-guide RNAs (sgRNAs) were designed by GPP sgRNA Designer ^9^ and synthetic oligos with chemical modifications (2’-O-Methyl at 3 first and last bases, 3’ phosphorothioate bonds between first 3 and last 2 bases) were synthesized by Synthego. Different combinations of sgRNAs targeting two ends of IG-DMR region were tested in 911 human cells to select sgRNAs with the highest cutting efficacy. Selected sgRNAs were then co-transfected with PX458 plasmid (Addgene #48138), which contains the expression cassettes of *S. pyogenes* Cas9 and GFP, into hiPSCs by Lipofectamine 3000 (Thermo Fisher Scientific) according to manufacturer instruction. After 24 hours post-transfection, single GFP+ cells were sorted by Astrios Cell Sorter (Beckman Coulter) directly in pre-warmed mTeSR medium supplemented with 10% CloneR (Stemcell technologies, Reference 5888) and hES Cell Cloning & Recovery Supplement (Stemgent, Reference 01-0014-500, dilution 1/2000). The single clones were kept at 37°C for 7 days, and the medium was changed every 2-3 days. Subsequently, genomic DNA of hiPSC clones was extracted by QuickExtract DNA Extraction Solution (Epicentre) and subjected to three PCR reactions with three sets of primers to screen for bi-allelic IG-DMR deletion (IG-DMR^−/−^). The positions of primers and the sizes of the PCR amplicons were illustrated in **Supplemental Figure 5A**. The sequences of selected sgRNAs and PCR primers for screening were detailed in **Supplemental Table 1-2**. We selected two single clones with no IG-DMR deletion in both alleles from the same screening process, used as control (IG-DMR^+/+^) in subsequent experiments. Off-targets of 2 sgRNAs used in this study were identified by CHOPCHOP ^10^ and detailed in **Supplemental Table 2-3**. The absence of any off-target in all clones used in this study were validated by Sanger sequencing (Genewiz).

#### Immunohistofluorescence and histochemistry staining

The transversal sections of 8-μm thickness were prepared from frozen muscles in the cryostat (Leica). The sections were subsequently fixed in 4% methanol-free paraformaldehyde for 10 minutes before blocked in the Blocking Buffer 10% Goat Serum (Gibco) in 1X PBS for 1 hour at room temperature. The blocked tissues were then incubated with primary antibodies in 1% Goat Serum (Gibco) in 1X PBS overnight at 4 °C. Samples were washed and incubated with Alexa-conjugated secondary antibodies (Life Technologies, dilution: 1/1000) in 1% Goat Serum (Gibco) in 1X PBS for 45 min at room temperature. After rewashing in 1X PBS, samples were mounted in DAPI Fluoromount-G (Southern Biotech) and stored at 4 °C. For enzyme histochemistry staining of succinate dehydrogenase (SDH) and Cytochrome C oxidase (COX) activities, freshly prepared slides were used and stained by using commercial kits (Bio-Optica, Reference 30114LY and 30115LY, respectively) according to the manufacturer’s protocols. Images were acquired with confocal microscopy TCS-SP8 (Leica) or AxioScan Slide Scanner (Zeiss).

#### *In situ* hybridization of miRNAs in skeletal muscle

For miRNA *in situ* hybridization (ISH), muscles were dissected and immediately fixed in 10% neutral-buffed formalin overnight before embedded in paraffin blocks. Transversal sections of 4-μm thickness were obtained in the Microtome (Leica). The ISH of miR-127-3p and miR-379-5p were performed by miRCURY LNA miRNA ISH Optimization Kit (FFPE) 1 (Qiagen, Reference 339451), according to the manufacturer’s protocol. All miRNA probes are double-digoxigenin-labeled and hybridized at a final concentration of 40 nM at 53°C for 1 hour. Probe with scramble sequence was used as the negative control. Nuclei were counterstained with Nuclear Fast Red (Fluka, Reference 60700). The miRNA probe sequences are shown in **Supplemental Table 1**.

#### Western blotting

Proteins were extracted from tissues or cells by RIPA buffer (Thermo Scientific) supplemented with Protease Inhibitor Cocktails (Complete PIC, Roche) and Benzonase 1:1000 (Millipore). Total protein concentration was measured by using Pierce BCA Protein Assay (Thermo Fisher Scientific). Equal amounts of protein were then loaded and separated by precast 4-12% Bis-Tris polyacrylamide gel (Thermo Fisher Scientific). Subsequently, the protein was transferred to a nitrocellulose membrane with the iBlot2 Dry Blotting system (Thermo Fisher Scientific). For detecting the proteins of interest, the membrane was blocked in Odyssey® Blocking Buffer (LI-COR) for 1 hour at room temperature before incubated with primary antibodies diluted in 50% Odyssey® Blocking Buffer overnight at 4 °C. After washed in 1X PBST, the membrane was incubated with secondary antibodies 1/5000 diluted in the 50% Odyssey® Blocking Buffer for 1 hour at room temperature. The membrane was then rewashed and blotting signals were acquired in the Odyssey Infrared Imaging system. Total protein (for muscle samples) measured by Revert 700 Total Protein (LI-COR) (for muscle samples) or Actin (for *in vitro* cells) was used as loading controls for quantification. Details of antibodies used in this study are presented in **Supplemental Table 1.**

#### Measurement of Citrate Synthase Activities

Citrate Synthase (CS) activity of muscle extracts was determined by Citrate Synthase Assay Kit (Sigma-Aldrich, Reference CS0720) according to the manufacturer’s protocol. 5-10 mg of muscle tissues were initially used and 5 μg of protein extract was used to measure CS activity.

#### Measurement of ATP concentration

ATP concentration of the muscle biopsies was determined by ATP Assay Kit (Abcam, Reference ab83355) according to the manufacturer’s protocol. 5-10 mg of muscle tissues were used in the assay. The precise mass of muscle tissues was noted and subsequently used to normalize across samples.

#### Measurement of mitochondrial and viral DNA

Total DNA was extracted from 5-10 mg of muscle tissues by using QIAamp DNA Mini Kit (Qiagen, Reference 51304) according to the manufacturer’s protocol. Purified DNA was subjected to qPCR using SYBR Green PCR assays (Thermo Fisher Scientific). Levels of genomic DNA were used to normalize data across samples. Primers amplifying genomic DNA, mitochondrial DNA and DNA from AAV vectors were detailed in **Supplemental Table 1**.

#### Measurement of mitochondrial OxPhos enzymatic activities

Enzymatic activities of complex I (NADH:ubiquinone oxidoreductase, rotenone-sensitive activity), II (succinate dehydrogenase, malonate-sensitive activity), III (decylubiquinol cytochrome c oxidoreductase, antimycin A - sensitive activity), IV (cytochrome c oxidase, KCN-sensitive activity); combined activities of complex I+III (NADH cytochrome c oxidoreductase, rotenone-sensitive activity) and II+III (succinate cytochrome c reductase, malonate-sensitive activity); and citrate synthase (CS) was determined as described ^19^. On the other hand, complex V (ATP synthase, oligomycin-sensitive activity) activity was measured according to Benit et al., 2006 ^20^. The measurement was performed in muscle homogenates for skeletal muscles and isolated mitochondria for hiPSC-derived myotubes. Muscle homogenates and myotube mitochondria were subjected to three freeze-thaw cycles to increase rotenone sensitivity. Subsequently, the OxPhos and CS activities were determined spectrophotometrically with Cary 60 UV-Vis spectrometer (Agilent) or Spark Cyto (Tecan). CS activities were used to normalize across samples as an indicator of mitochondrial mass.

